# *Anopheles coluzzii* infection by the microsporidian, *Vavraia culicis*: the effect of host age

**DOI:** 10.1101/2020.10.31.363077

**Authors:** Nayna Vyas-Patel

## Abstract

Host age at infection has important implications for disease development. In mosquitoes, infections with microsporidia and later concurrent infections with malaria parasites, leads to a suppression in the development of malaria parasites. Host age at infection with microsporidia could have implications for disease outcomes when infection occurs subsequently with malaria parasites. Mosquito larvae can take between five to seven days or more depending on the temperature to reach the adult stage, giving the microsporidian *Vavraia culicis*, a theoretical head start in establishing and developing within larvae and possibly resulting in different levels of infection in emergent adult mosquitoes. To determine the effects of early or late infection with *V. culicis*, equal numbers of *Anopheles coluzzii* larvae were infected individually with a high or low dose of *V. culicis*, at different ages post hatching.

Significantly fewer spores were produced from mosquitoes infected later, than ones infected earlier with microsporidia and there was an initial delay in the production of spores from later infected mosquitoes. In early infected larvae, there was no such initial delay and spore production took off unchecked. The infectious dose of *V. culicis* did not affect the total spore count per mosquito. Male mosquitoes produced fewer spores than females. Daily mosquito longevity and pupation was not affected significantly by infection, the infectious dose of *V*. *culicis* given or by the sex of the mosquito. Considering hourly deaths, early infected hosts died 17 to 18 hours earlier than later infected larvae. The number of *V. culicis* spores rose with increasing duration of infection. When equal duration of infection was considered, the findings remained the same. Host age at infection influences disease outcomes and virulence.

## Introduction

*Vavraia culicis* (Weiser J 1947 & 1977; Vávra J and Becnel JJ 2007), a microsporidian parasite with a wide host range, naturally parasitizes a number of mosquito genera and species (Vávra J & Becnel JJ 1977). This obligate and horizontally transmitted parasite has been the subject of numerous studies, not merely to explore its potential as a biological control agent (Kelly JF et al, 1981; Reynolds DG 1970; Agnew P et al, 1999), but also for its capacity to interfere with the subsequent infection and establishment of *Plasmodium* species (malaria parasites) in the adult mosquito (Bano L 1958, Bargielowski I & Koella JC 2009; Lorenz LM & Koella JC, 2011).

Mosquitoes have life stages in two different mediums – larvae develop to pupation in water and emerge as flying adults into the terrestrial world. Each stage (larval and adult) has its own developmental age and challenge. Mosquito larvae first encounter the infective spores of the microsporidian *V. culicis* in the aquatic environment, however, nature is not always uniform and at any given time in a tropical environment, a typical mosquito breeding pool generally harbours mosquito larvae of varying ages and species (Karuitha M, 2019 & Vyas-Patel N, 1988), some of which could have been infected with parasites if present and others which had yet to be infected. A mix of larval ages and indeed species could be present in a mosquito breeding pool; seldom will a pool contain larvae of uniform age as it would continuously be used by different adult mosquito species to lay eggs at different times. The distribution of spores in water in the field is not known, it might be patchy (clumped) or evenly (uniformly) distributed in water affecting spore uptake by mosquito larvae. Little is known about the effect of larval host age on the development of the parasite, especially as mosquito larvae are infected horizontally with *V. culicis* in water (spores are ingested) and this can happen at any larval age.

The virulence of *V. culicis* infection at different host ages was also of interest. Virulence being the degree of pathogenicity of a parasite and determined by its ability to invade and multiply within a host. Does pathogenicity, hence parasite virulence, increase with increasing host age at infection in *V. culicis*? Futuyma DJ (2001) stated that the larger the parasite load, the higher the virulence of the parasite and the greater its potential to cause harm and consequently, host fatalities. Therefore, it was postulated that the larger the numbers of *V. culicis*, the greater its virulence and capability to cause harm to the host, hence the numbers of *V. culicis* spores at the end of a mosquito’s life was a key indicator of its virulence. If these numbers differed, it would give some indication of *V. culicis* virulence within younger and older infected mosquitoes.

The trade-off between parasite virulence and its survival and the concept of optimal virulence have been elucidated for many host parasite interactions. A number of factors affect the virulence of *V. culicis* in mosquitoes. Intraspecific competition, host parasite interactions and host condition were explored by Michalakis Y et al (2008) as factors affecting *V. culicis* virulence. Using four different isolates of *V. culicis*, Bargielowski I and Koella JC (2009) found that three of the isolates were more virulent than others and gave rise to higher spore counts at the end of a mosquito’s life, hence virulence can change between different isolates of microsporidians. Bedhomme S et al, (2004) demonstrated that consuming host resources is another factor that influences a parasite’s virulence and that the nutritional status of the host was an important factor. As older hosts provide larger reserves of food (being larger in size), it would be natural to conclude that infections of older hosts should result in larger numbers of spores, a factor explored here.

Studies have examined the effect of host age on parasitism by microsporidia in different insect species. In the red flour beetle *Tribolium castaneum*, investigators noted that virulence to the host and the success of its microsporidian parasite *Nosema whitei* was mainly determined by host age at infection. Furthermore, it was found that infection was only possible in young larvae and that the disease prolonged the life span of the larval stages and prevented them from developing to pupation (Blaser M & Schmid-Hempel P, 2005). Similarly, different workers investigating *N. whitei* in *T. castaneum*, concluded that age structure, spore concentration, and timing of infection (i.e. host age when infected) were important factors to consider in the reduction of infestation of flour by the beetle (Onstad DW and Maddox JV, 1990). In Monarch butterflies, early instars were more susceptible to a higher spore dose of their microsporidian parasite, *Ophryocystis elektroscirrha* than when the later, older larvae were infected, although all larval stages tolerated the parasite at the lower doses. At the highest spore dose, the early instars decreased survival to eclosion (Altizer SM & Oberhauser KS 1999). Vijendravarma RK et al, (2008), found that the susceptibility of *Drosophila melanogaster* by the microsporidian *Tubulinosema kingi*, decreased with larval age, only the early stage larvae were highly susceptible and that the microsporidian only proliferated to a small degree in the larval stages and increased replication greatly within the adult stages. Considering larval age in mosquitoes when infected with parasites other than microsporidia, Umphlett CJ and Huang CS (1973) found that early instars of *Aedes aegypti* infected at a lower dose with the fungus *Lagenidium*, were highly susceptible to the fungus compared to older larvae. Similarly, when larvae of *Aedes aegypti* were infected with the mermithid nematode, *Romanomermis culicivorax* early instars were more susceptible than when later, fourth instars or pupae were infected (Vyas-Patel N, 1983). These studies indicate that larval age at the time of infection by microsporidia and by other parasites is a factor that influences outcomes for both the parasite and host. *Vavraia culicis* is considered a useful agent in the suppression of *Plasmodium* species within adult mosquitoes (Bargielowski I & Koella JC 2009), it is essential therefore, to ascertain any impact of larval host infections at different ages on adult parasite load. Clearly, if the microsporidian spore load is low or high in an emergent adult due to differential larval infection, this could affect subsequent infection and interference with later infections of the adult with malaria parasites.

To examine the effect of larval age, two groups of *Anopheles coluzzii* larvae were infected with *V. culicis* spores on day one, then day three, five and seven after hatching. Group one larvae were infected at a low dose of 5,000 spores per individual mosquito and group two infected at the higher dose of 50,000 spores per mosquito. All groups including controls, received the optimum daily food dose and they were reared under ideal insectary conditions. The day of pupation and death was noted for every mosquito. As only females carry and transmit the malaria parasite, wings were removed and measured from the experimental and control female mosquitoes; wing lengths being highly correlated with mosquito weight (Lounibos LP et al, 1995). Spores were counted from every mosquito infected with *V. culicis*, after the death of the adult. All of the mosquitoes were given the same nutrients and conditions and were allowed to die naturally, without any intervention at any stage.

## Materials and Method

*Anopheles gambiae* N’guesso strain, reared at Imperial College London (ICL) and subsequently at the Silwood Park, Ascot, campus, were used. Maintained over many years at ICL, this colony known to be from Yaoundé, Cameroon was reclassified as *Anopheles coluzzii* (previously *An. gambiae* ‘M’ form) following molecular characterisation (Habtewold T et al, 2016). Note that previous publications (prior to 2016) referring to *Anopheles gambiae* from Yaoundé Cameroon and reared at ICL, are more than likely to be the *An. gambiae* ‘M’ form strain, i.e. *Anopheles coluzzii*.

Mosquito larvae were infected at different ages; day one, three, five and seven days post hatching, with *V. culicis* and compared to uninfected mosquito larvae. Two different infection levels were used, group one was infected with the lower dose of 5,000 spores per mosquito larva and group two with the higher dose of 50,000 spores per larva. All groups and controls received optimum nutrition and rearing conditions. The day of pupation was noted for every larva as was the day of adult death. All of the female mosquito wings were measured and a spore count carried out for every infected mosquito used in the experiment.

### Experimental Design

A total of 1,800 newly hatched mosquito larvae were individually placed in a 12 well plate; one larva per well, in 2 ml of deionised water per well. Falcon Multiwell™ 12 well plates, (Becton Dickinson) were used.

The first row of four wells in a plate was allocated to group one larvae (low infections), the second row was allocated to group two larvae (high infections) and the final four rows were allocated to the control larvae which were not infected with *V. culicis*. For subsequent plates, treatment rows were allocated randomly. This ensured that all the treatments were included in every individual plate and the rows were allocated randomly. A fully random allocation of treatments (with every well selected at random for treatment) could have been used, but was avoided; as such an attempt had previously resulted in pipetting errors. This way, accuracy could be maintained along the rows; without compromising random allocation of treatments.

### Larval Food Regime

TetraMin^R^ fish food was weighed and made up to 300ml in a beaker, so that when 2 µl of the food was pipetted into the wells, the larva in each well received 0.05mg of food on day one, then 0.06mg on day two, 0.1mg on day three, 0.16mg on day five, 0.32 mg on day six, and 0.6 mg on subsequent days. This gentle increase in food dissemination ensured that mosquito larvae received the optimum food levels for their size, without the attendant growth in bacteria that would occur if too much food was placed into the wells. The food was freshly prepared daily and the beaker placed on a magnetic stirrer to ensure even distribution of the food particles, whilst 2µl aliquots of the food solution were applied to each well.

### Mosquito Rearing

*Anopheles coluzzii*, originating from Yaoundé Cameroon and maintained over many generations in the laboratory at Imperial College London (ICL), was used for the experiment. Mosquitoes were reared in a sealed insectary, kept at a temperature of 26°C (+/-1), with a 12:12 hour light: dark cycle and a humidity of 70% (+/-5). Adults were blood fed on a human arm introduced into the rearing cage and allowed access to sugar in the form of a sugar cube placed inside the cage. Eggs were collected from a shallow container, lined with filter paper around the sides, half filled with water and placed inside the cage.

### Experimental Mosquitoes

Once an experimental larva had pupated in a well, the day of pupation was noted and the pupa transferred to an Eppendorf tube, placed lid open, in a 10 ml Falcon tube. Mosquito netting was secured to the open end of the Falcon tube and cotton wool soaked with a 10% sucrose solution placed on top of the netting. This ensured that when adults emerged (usually within 24 hours), they had unlimited access to sucrose. Thus, each mosquito was reared individually from larva to adult.

Falcon tubes were examined daily for adult death and fresh sucrose supplied. The day of death was noted and the mosquito was transferred to a labelled Eppendorf tube and stored in the fridge. The sex of each adult was noted. Female wings were removed for measurement from every adult female (whether infected or not). Finally, all the infected adults were individually crushed in their Eppendorf tube with a 1.5ml plastic pestle and the spores counted using a haemocytometer.

### Microsporidian Cultures

*Vavraia culicis floridensis* spores, initially obtained from a stock culture maintained by J.J. Becnel (USDA Gainesville, USA) were cultured by infecting batches of 40, two day old mosquito larvae in Petri dishes with a dose of 40,000 spores per 40 larvae. The spores were counted using a haemocytometer, under a phase contrast microscope. The infected larvae were reared to pupae in the Petri dishes and on pupation, were transferred to cages for adult emergence. Adults were collected after two weeks, placed in a fridge and collections of three or four were crushed in 1 ml of deionised water in an Eppendorf tube with a 1.5ml plastic pestle to release the spores. The culture was pooled and aliquots of it counted and made up to the required dilutions, as and when required for experiments. For this experiment, a large stock culture was counted and diluted into two flasks to ensure that 250<u>µ</u>l aliquots from one of the flasks would contain 5,000 spores and 250 aliquots from the second flask would contain 50,000 spores. It was ensured that the volume of the dilution in each flask was enough to apply to all 600 larvae in both sets.

### Wing Length Measurements

After female mosquitoes had died, both wings were detached with forceps and placed on a drop of water on a microscope slide and covered with transparent Sellotape. A row of labelled wings was prepared on a slide. The slides were scanned alongside a ruler on a scanner attached to a computer. The scanned images of the wings were measured using the program Image J, (http://rsb.info.nih.gov/ij/).

### Statistical Analysis

Statistical analysis was performed using R (http://www.r-project.org/) and Windows Excel. A maximal quasibinomial Generalised Linear Model (GLM) was fitted to the data; this contained all factors, interactions and covariates that might be of any interest (maximum likelihood estimation). This would indicate the minimal adequate model to describe the data. Spore count was the response variable; day infected, sex of the mosquito, infection level were the explanatory variables. Model simplification was carried out by stepwise elimination of non-significant factors and interactions.

## Results

The raw results together with the computation from ‘R’ statistical analysis are presented in supplementary materials as Excel and .text files.

A maximal model was fitted to the data and the significance of terms was tested by stepwise deletion. The data were over dispersed and the appropriate model was a quasipoisson Generalised Linear Model (GLM). Spore count was the response variable, day infected, sex, infection level were the explanatory variables.

It was found that there were no significant interactions between the explanatory variables (i.e. between day infected, sex of the mosquito, female wing length and infection level). The original level of infection did not affect the spore count at the end of the mosquito’s life (t=0.501, d.f. =1, p=0.62), i.e. the infectious dose did not affect spore count. Male mosquitoes produced fewer spores than did females (t=3.31, d.f.=1, p<0.01). The day infected affected spore production; later infections produced fewer spores (t=9.45, d.f. =1, p<0.001) (Table 1, Figure 1,).

**Table 1.**
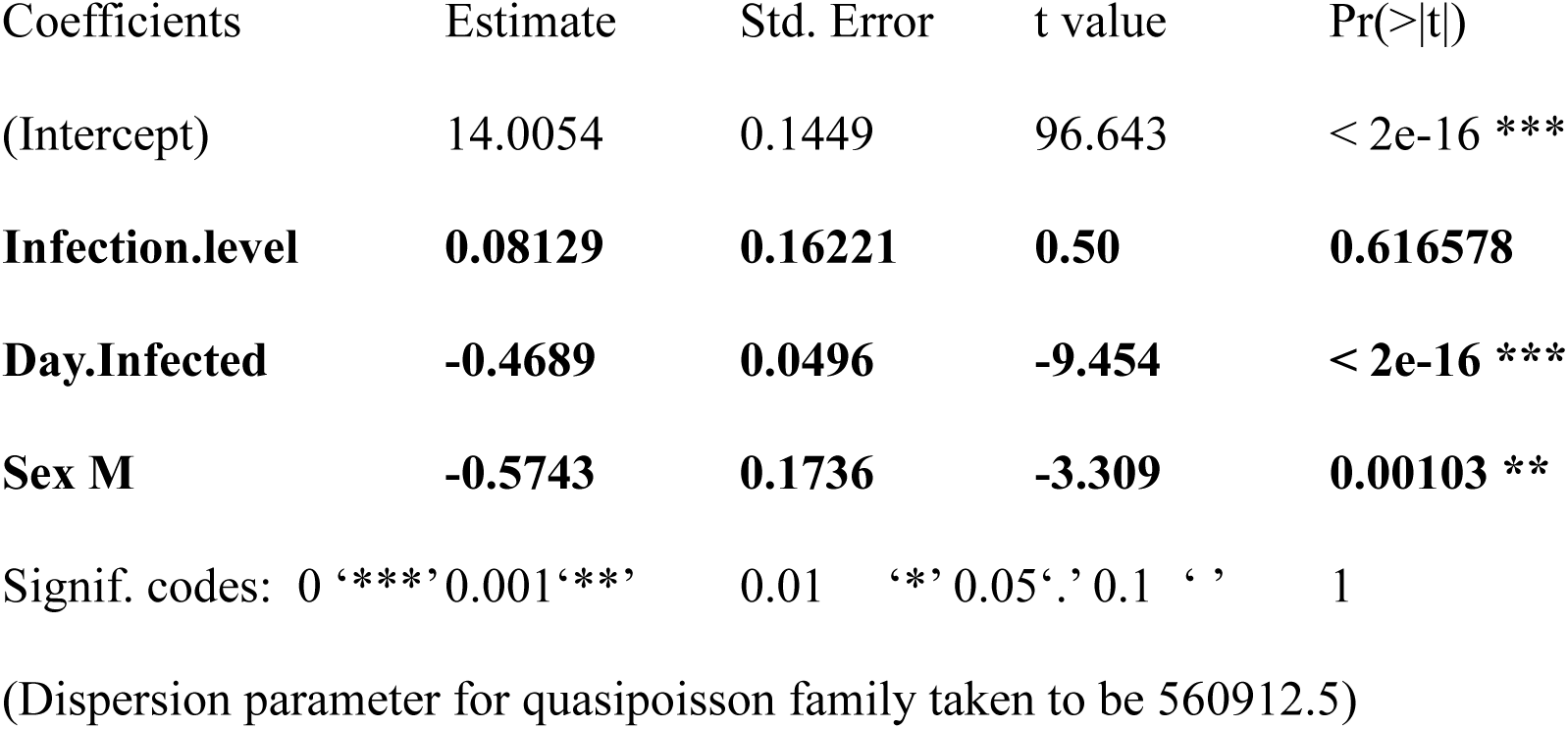
No significant interactions between the explanatory variables - i.e. between day infected, sex of the mosquito, female wing length and infection level.

**Figure 1:**
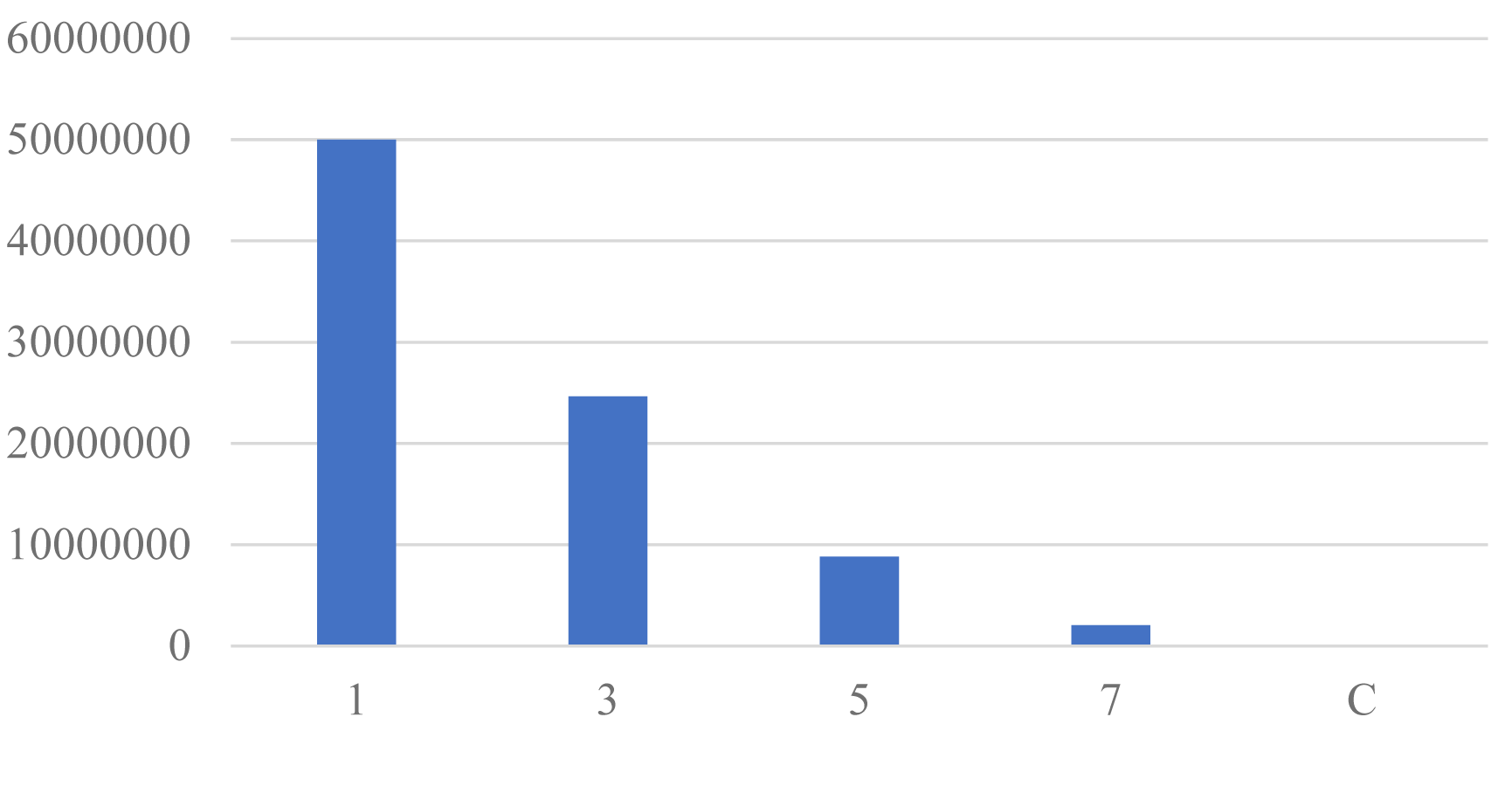
Sum of Spore.count (y axis) by Day.Infected (x axis) Mosquitoes infected later (older larvae) resulted in fewer spores in the adults compared to early infected larvae.

Infection with *V. culicis* does not affect the age at death of the mosquitoes (t = 0.77, d.f. = 598, p-value = 0.44), Table 2.

**Table 2.** Infection does not affect the age at death of adult mosquitoes.

| Coefficients: | Estimate | Std. Error | t value | Pr(>\|t\|) |
| --- | --- | --- | --- | --- |
| (Intercept) | 20.2950 | 0.4959 | 40.924 | <2e-16 *** |
| **Infected** | **-0.4700** | **0.6074** | **-0.774** | **0.439** |

Signif. codes: 0 '\*\*\*' 0.001 '\*\*' 0.01 '\*' 0.05 '.' 0.1 ' ' 1 .Residual standard error: 7.013 on 598 degrees of freedom. Multiple R-squared: 0.001, Adjusted R-squared: -0.0006702; F-statistic: 0.5988 on 1 and 598 DF, p-value: 0.4393.

In order that the same time frame available for spore production could be considered in both early (younger larvae) and late (older larvae) infected hosts, the duration of infection was considered. The day of infection was taken away from the day of mosquito death and the spore count for the same duration was considered. When duration of infection was considered (day died – day infected = duration of infection), it did not affect the results. Once again it was seen that the original level of infection, i.e. whether larvae were infected at a high or low dose, did not affect the number of spores at death (t=0.12, d.f. = 1, p = 0.9). As expected, the number of spores rose with the duration of infection (t=8.9, d.f. =1, p<0.001). Furthermore, when duration of infection was considered, the age of the mosquito at infection affected the spore count at death, as older mosquitoes not only produced fewer spores, but were initially able to delay the onset of spore production (t=5.9, d.f. = 1, p<0.001), Table 3 and Figure 5.

**Table 3.**
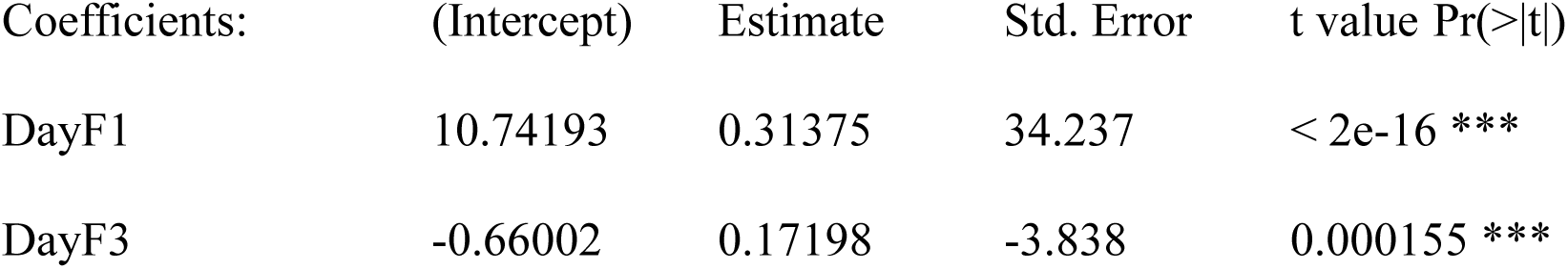
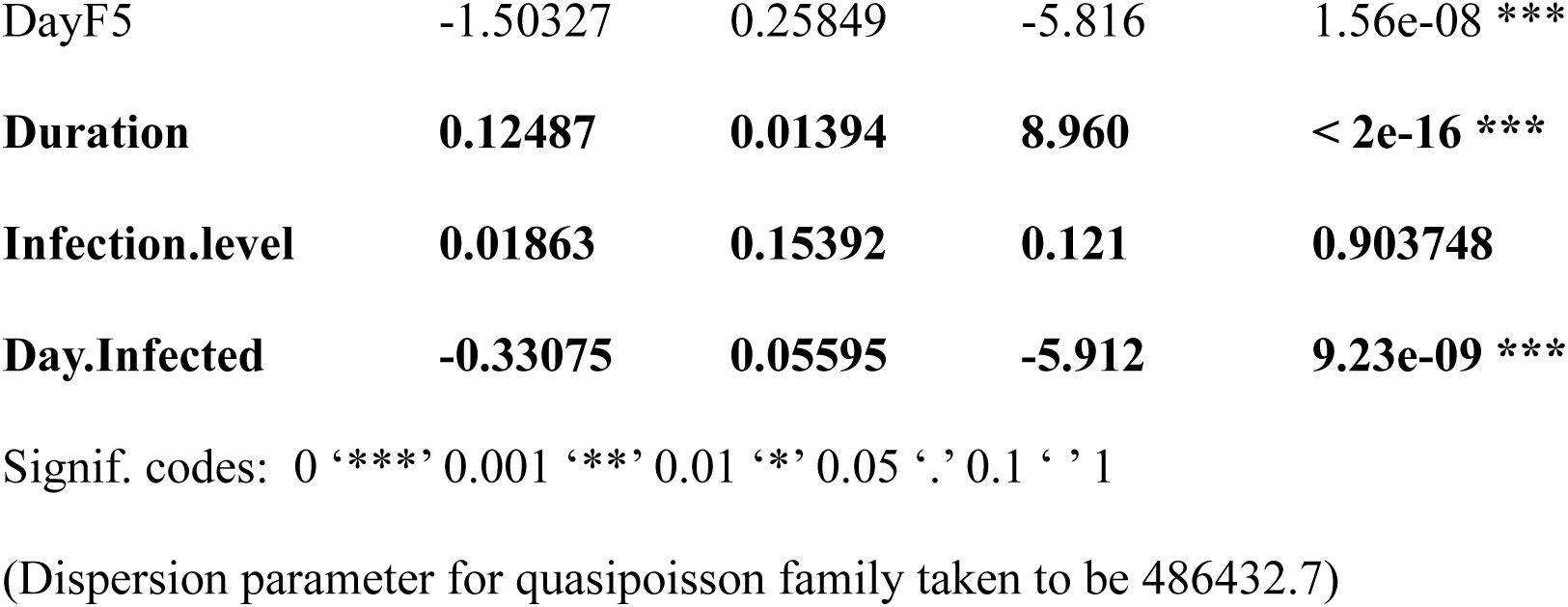

| Coefficients: | (Intercept) | Estimate | Std. Error | t value | Pr(>\|t\|) |
| --- | --- | --- | --- | --- | --- |
| DayF1 | 10.74193 | 0.31375 | 34.237 | < 2e-16 | *** |
| DayF3 | -0.66002 | 0.17198 | -3.838 | 0.000155 | *** |

|  |  |  |  |  |
| --- | --- | --- | --- | --- |
| DayF5 | -1.50327 | 0.25849 | -5.816 | 1.56e-08 *** |
| **Duration** | **0.12487** | **0.01394** | **8.960** | **< 2e-16 \*\*\*** |
| **Infection.level** | **0.01863** | **0.15392** | **0.121** | **0.903748** |
| **Day.Infected** | **-0.33075** | **0.05595** | **-5.912** | **9.23e-09 \*\*\*** |

Signif. codes: 0 '\*\*\*' 0.001 '\*\*' 0.01 '\*' 0.05 '.' 0.1 ' ' 1
(Dispersion parameter for quasipoisson family taken to be 486432.7)

Figure 5 indicates that the rate of growth of *V. culicis* does not differ with the age at infection (the slope is the same), but the numbers of spores were depressed initially with later infection (the intercepts are lower with greater age at infection). Between days 10 and 20, spore production appeared to reach a maximum plateau in all cases. As before, the longer the duration of infection (early infected larvae), the higher the spore count.

The age at pupation was not affected by the initial dose of *V. culicis*, or by the sex of the mosquito, however there was a small delay in the onset of pupation with later infection, Table 4.

**Table 4.**
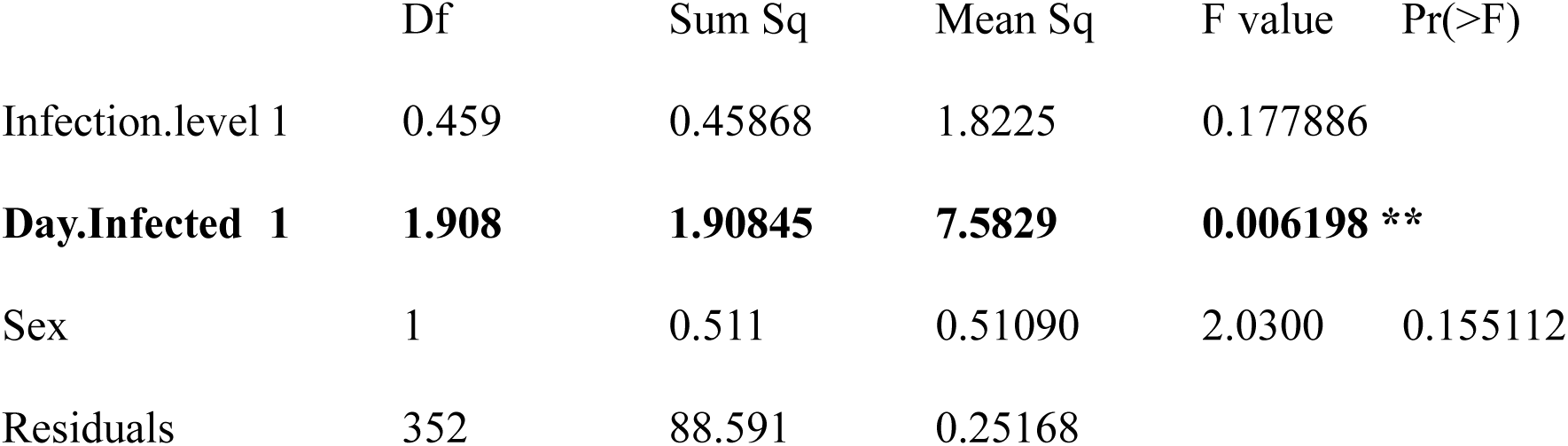

The above Anova indicated that there was an effect on pupation, to resolve the delay, a linear model was fitted with the only significant term in it (day infected). This indicated that early infected mosquitoes started to pupate on day 7 (Table 4), but that the onset of this pupation was delayed by 0.03 days (about an hour and a half) for every day later they are infected (Table 5). Whilst this is statistically significant, given that pupation was noted only once a day, it is not meaningful to infer that pupation was affected in later infected hosts, Table 5.

**Table 5.**
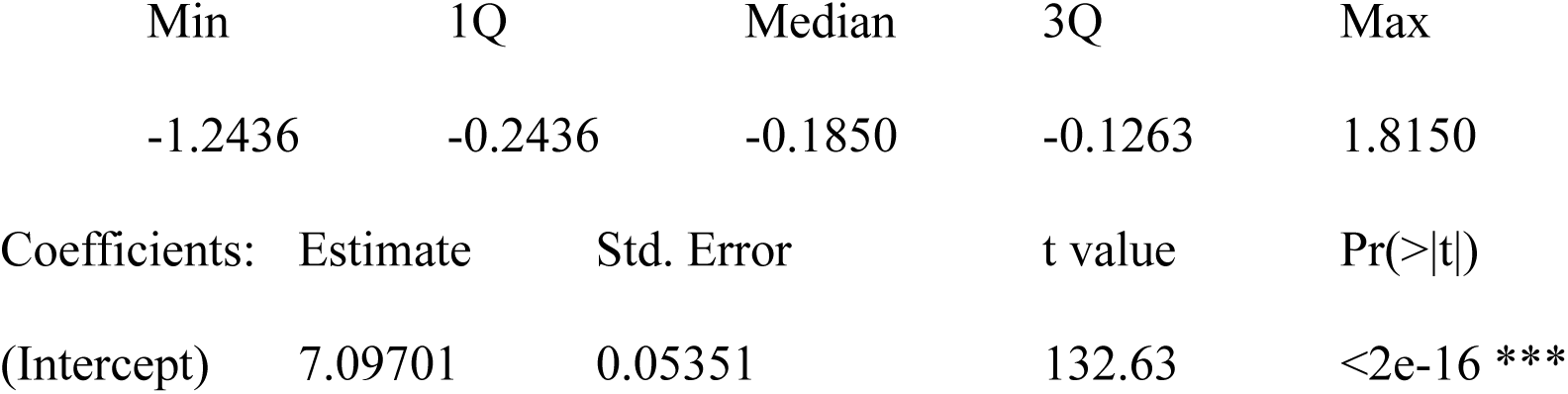
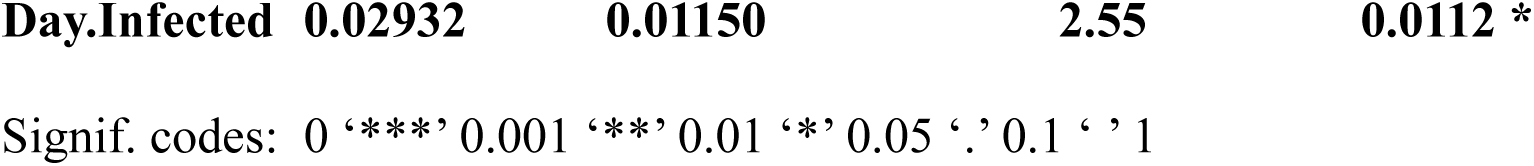

Examining the age at which mosquitoes died, the only important effect was the time of infection (larval age). The interaction between the level of infection and the day they were infected did not appear to matter, Table 6.

**Table 6.**
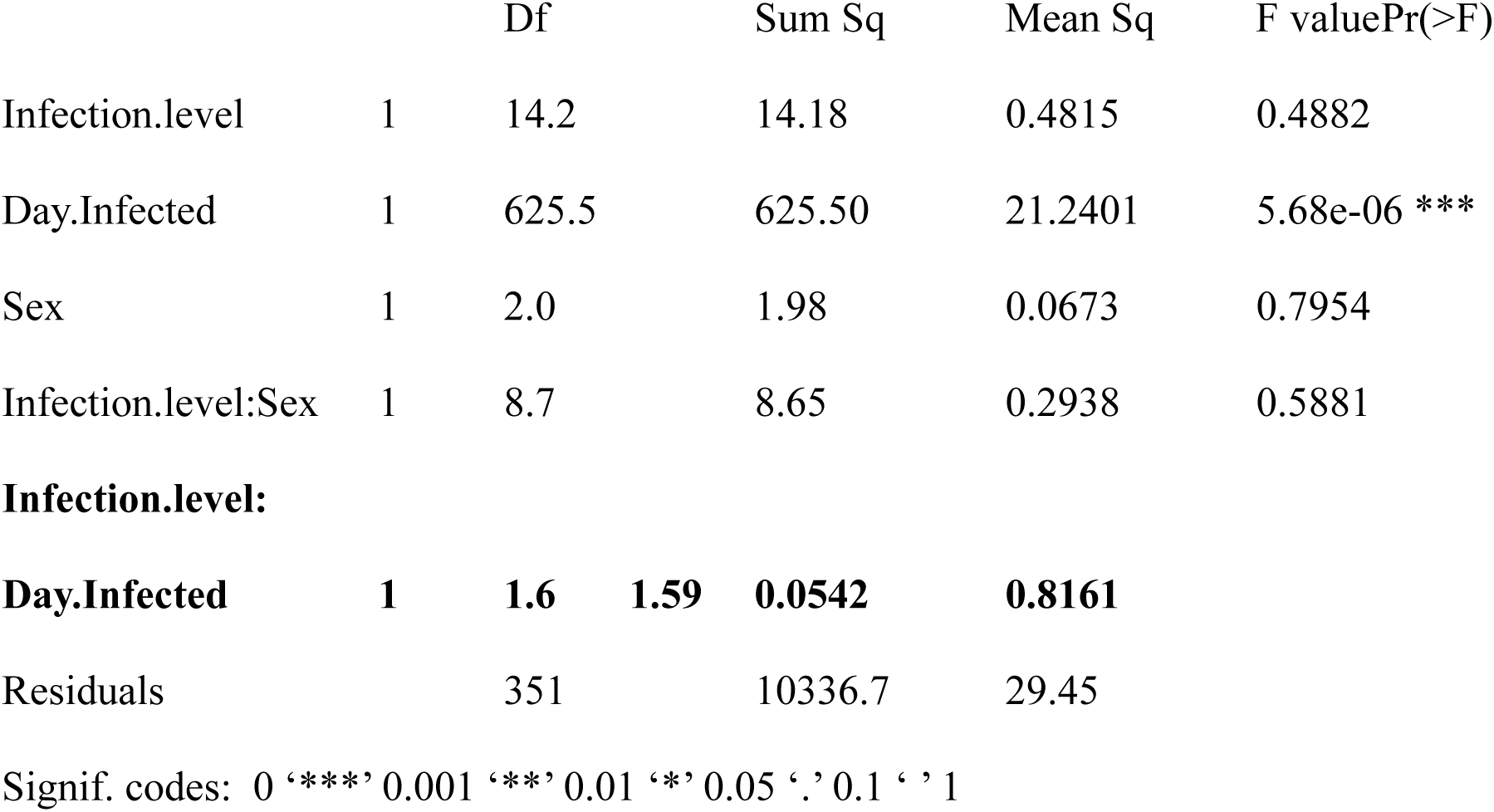

The minimal adequate model only has the day infected in it and a linear model fit suggests that early infected mosquitoes died earlier than later infected mosquitoes (about ¾ of a day (17-18 hours) of extra life for every day later infected), Table 7.

**Table 7.**
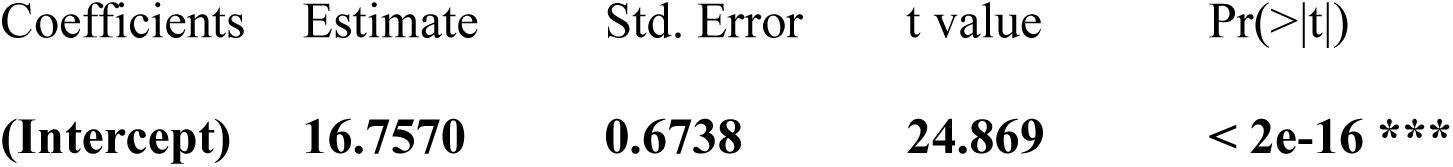
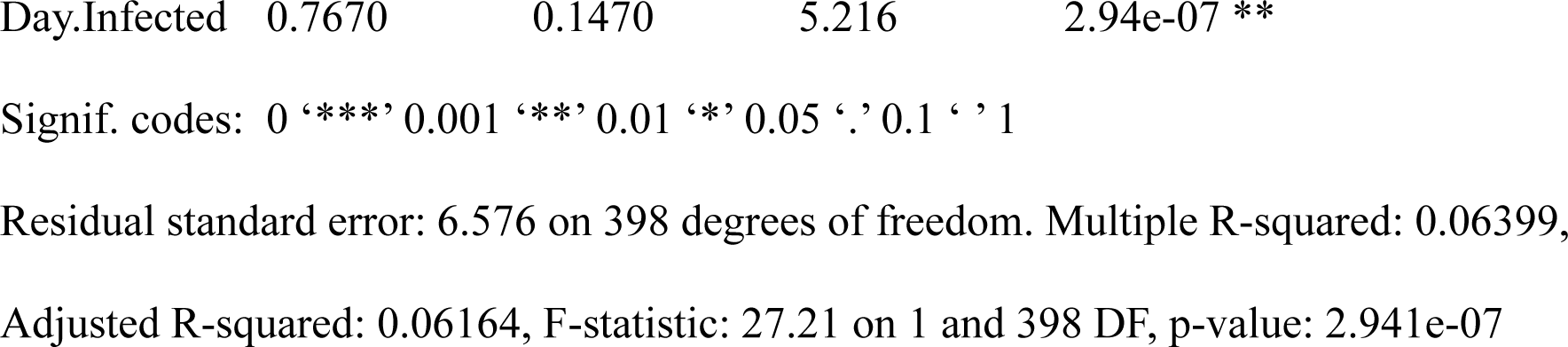

Examining adult longevity of the mosquitoes, it was found that neither the sex of the mosquito, nor the initial dose of *V. culicis*, mattered, Table 8.

**Table 8.**
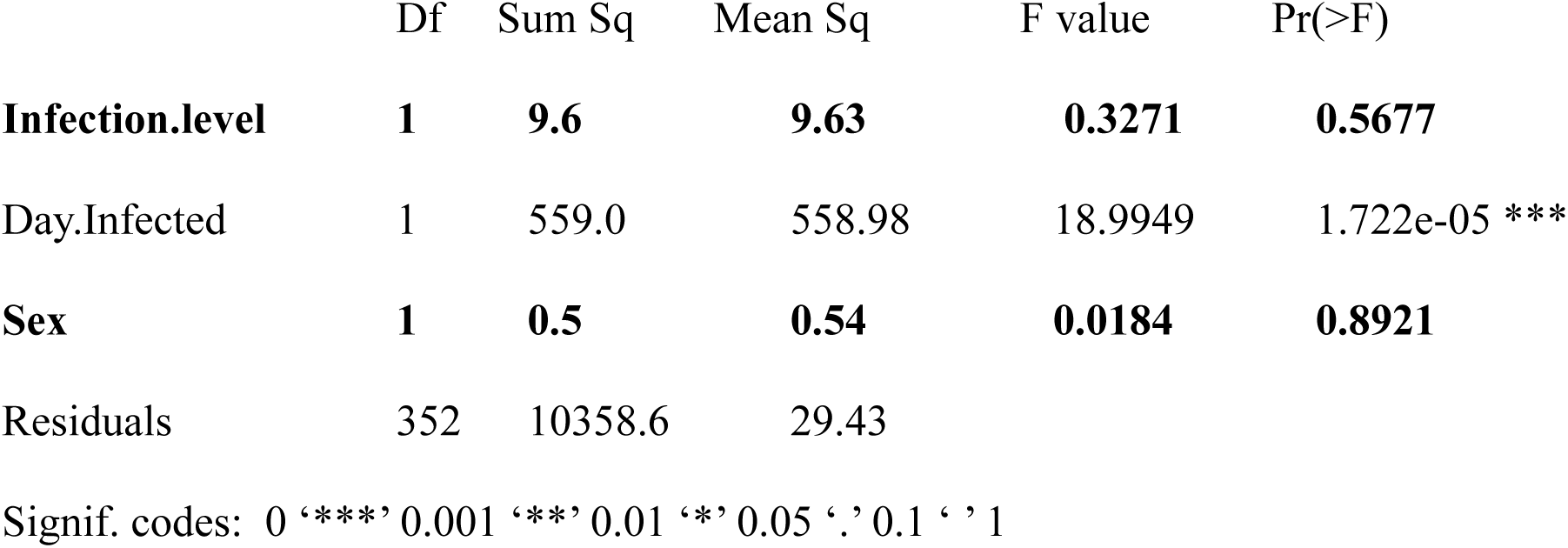
Analysis of Variance Table Model 1: Ad ∼ Infection.level * Day.Infected * Sex Model 2: Ad ∼ Infection.level + Day.Infected + Sex> summary(ad2)

Using adult longevity data, once again the minimally adequate model indicated that the earlier mosquitoes are infected, the earlier they die (about ½ a day longer) for each delay in infection, Table 9. This is similar to the results seen in Table 7.

**Table 9.**
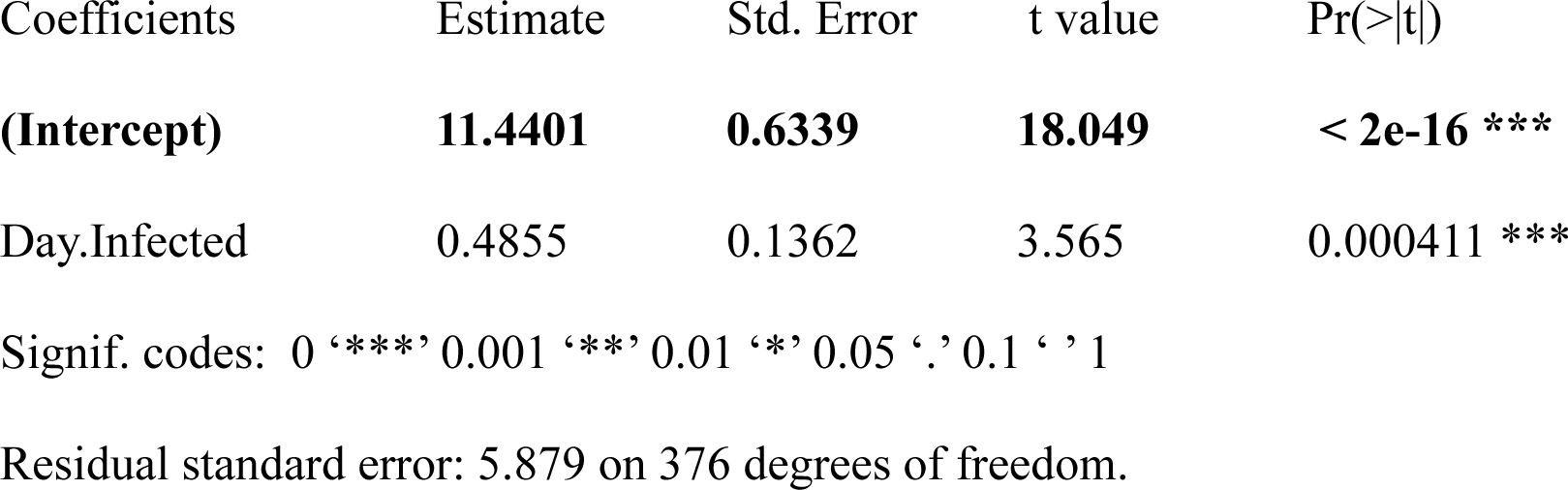

Analysing the wing measurement data, it was found that there were no significant interactions between wing length and spore count, dose of infection, or mosquito age at infection.

## Discussion

A difference exists in spore production in larvae that were infected at a younger, compared to an older age. Later infected, older larvae, produced significantly lower numbers of spores compared to early infected, younger larvae (Table 1, Figure 1). This effect persisted even when the duration of infection was considered, that is when comparing the same duration in early (young) and late (older) infected larval groups. There was an initial delay in spore production from older infected hosts (Figure 5). This initial delay may account for the lower spore counts from older, compared to younger infected larvae.

It is known that the aquatic environment presents a greater challenge in terms of the density and diversity of microorganisms that can infect mosquito larvae compared to the adult aerial stage. League GP et al (2017) concluded that the high density and diversity of microorganisms encountered by larvae meant that selection pressure equipped surviving larvae with highly robust immune systems to ensure development to the adult stage.

Furthermore, that larvae possessed greater numbers of circulating haemocytes and stronger lytic and melanisation capacity than adults (League GP et al, 2017). Similarly, Brown LD et al (2019), demonstrated that mosquito larvae were more proficient than adults in killing bacteria, and that this correlated with the stronger cellular and humoral immune responses in larvae compared to adults. Clearly larvae are not born with a fully developed immune system, otherwise none of the spores ingested by one day and three day old larvae could develop; to the contrary, the opposite was found here, spore production rose unchecked from younger larvae. Examples from other microsporidian infections (Altizer SM and Oberhauser KS 1999,Blaser M & Schmid-Hempel P 2005, Milner RJ 1973, Maddox JV 1990 & 2000, Vijendravarma RK et al, 2008) and of fungal and nematode pathogens of mosquito larvae (Umphlett CJ and Huang CS, 1973; Vyas-Patel N, 1983) similarly indicated that younger, rather than older larvae were more susceptible to infection by parasites and pathogens in their environment. It is likely that younger larvae had yet to fully develop their immune system, in contrast to the fully developed immune systems of older larvae, resulting in younger larvae being more susceptible to infections. This could explain why, even when similar duration of infection was considered, older infected larvae with their fully developed, robust, immune system produced fewer spores than larvae infected at a younger age. It could also explain the initial delay in spore production from older hosts, as they should be better able to mount a stronger immune response than younger larvae whose defence response had yet to fully develop. The results seen here were not merely to do with the length of time the spores had to develop, more that the mature immune system of older larvae was more proficient at keeping pathogens at bay.

Parasite virulence from younger infected larvae appeared to be greater than from older infected hosts as larger numbers of spores were retrieved from younger infected larvae. The higher spore counts from early infected larvae indicates that virulence is higher in this group, regardless of their smaller body size and that the optimal food and environmental conditions provided, buffered early infected mosquitoes and ensured damage limitation.

Infection with *V. culicis* did not affect the daily age at death between the mosquito groups, Table 2 and Figure 4. Closer scrutiny revealed that early infected larvae died 17 to 18 hours earlier than later infected mosquitoes. Although this was just under a day earlier (Table 7), this difference could be greater in the field as the mosquitoes were reared individually under optimal conditions here. Furthermore, any time difference would be enough to alter malaria transmission in cases of subsequent infection of the host with malaria parasites, killing the mosquito before the average nine to ten days that malaria parasites require to mature and transmit the disease in the field. This is not to suggest that *V. culicis* should be advocated for use in the field with a view to suppressing the development of subsequent malaria parasite infections in mosquitoes, as *V. culicis* is closely related to *Trachipleistophora hominis*, an opportunistic microsporidian parasite known to infect both mosquitoes capable of transmitting malaria and other diseases and immunocompromised individuals, especially AIDS patients (Cheney SA et al, 2005). The aim here was to examine the factors affecting microsporidian infected hosts. Other microsporidians species are known, that are host specific and incapable of infecting humans. These species could be better biological control candidates (Andreadis TG, 2007) and might also achieve such an effect, i.e. the suppression of secondary infections, especially of malaria parasites.

Physical traits such as gut volume and content could also be factors of influence in spore production in larvae infected at an earlier age which were more susceptible to infection and proliferation by microsporidia and produced greater numbers of spores. It may be easier for microsporidia to multiply rapidly in the relatively immature cellular structure proffered by younger larvae compared to the more mature and densely packed body of older larvae. Gut volume (Weiser J, 1969), gut content and histological reorganisation in preparation for pupation (Milner RJ, 1973), may present contributing factors that affect spore production in older larvae.

Differences in larval behaviour between younger and older *Drosophila melanogastor* larvae was reported to be the reason for older infected larvae producing fewer spores of *Tubilinosema kingi* than younger infected larvae (Vijendravarma RK et al, 2008). However, a behavioural difference between younger and older larvae is unlikely to be the reason for the differences in spore production seen here. Most mosquito larvae use two feeding methods, they can filter feed from surface microorganisms, secondly, they can scrape biofilms from underwater vegetation and rocks (Roberts D, 2014). Both younger and older larvae had no option but to ingest the spores present in the experimental wells, along with their food particles. Ingestion of spores was inevitable and unavoidable, with little scope for any behavioural differences, at least with regard to spore ingestion.

In experiments considering the response of *Aedes aegypti* larvae to *V. culicis*, a study of the proteome of *Aedes aegypti* by Biron D (2005), came to the conclusion that older larvae reacted most strongly to infection (in terms of the percentage of protein spots whose expression was affected). In Biron’s 2005 study, the mosquitoes had all been infected at the same time with *V. culicis* and examined at different ages, here they were infected at different ages and spores counted at death. The stronger immune reaction from older larvae was attributed to the longer time the microsporidian spores had to mature in the host in Biron’s study (2005). The results here suggest that the immune response to infection differs in its ability to hamper spore production from younger and older larvae. Unchecked exponential increase in spore production was seen from younger infected hosts, whilst there was a delay in spore production initially from older hosts (Tables 4 & 5), which could be the result of a stronger immune response from older hosts and ultimately this had an impact on the levels of total spore production.

Early infected larvae died earlier than later infected (Table 7), hence control programmes using *V. culicis* aimed at reducing malaria transmission should ideally target younger larvae. However, as the difference in time of death between early and late infections was only a matter of hours, this should not be critical. More fundamental would be to know whether infecting older larvae brings about the same suppressive effect on the development of the malaria parasite, as found by Bargielowski I & Koella JC (2009) from early infections, where two day old larvae were infected and found to subsequently interfere with the development of malaria parasites in the adult.

Neither the sex of the mosquito nor a higher infectious dose affected the longevity of mosquitoes. The spore count on death was not affected by the infectious dose of spores used. Improved spore storage conditions over time needs to be investigated for *V. culicis* as this might improve spore viability. In general, all microsporidian spores deteriorate in UV light (Li X et al, 2003) hence the type of lights used in rearing rooms may affect the viability of spores. Spore counts were lower from males (Tables 4, 5 and 8). It is known from a previous study that the wings of males tested here were smaller than those of females (Vyas-Patel N, 2019). In a number of mosquito species, male mosquitoes have been found to be smaller in size than females (Thornhill R & Alcock J, 1983) and it would have been in keeping if a higher infectious dose led to males dying sooner than females. In the present study, female *An. coluzzii* wing lengths were not affected by any of the factors tested, indicating a degree of tolerance to microsporidian infection as found by other workers (Andreadis TG 2007) and an inability to clear the infection. That male mosquitoes produced fewer spores than females (Tables 4 & 5) was probably due to their smaller size limiting nutrient store space compared to the larger females as indicated by Agnew P et al, (1999). Infection with *V. culicis* depletes the food resources of infected mosquitoes and a lack of host nutrients is a limiting factor for spore development (Rivero A et al, 2007; Bedhomme S et al, 2004), so the smaller the food store, the fewer the spores. Studies of *Nosemi whitei* in *Tribolium castaneum* have similarly noted that spore count at death correlated with host body size (Blaser M & Schmid-Hempel P, 2005). Here, it could also be that as male mosquitoes do not have to reserve or divert their energy stores for the production of eggs, they may be able to invest more energy in trying to ward off the infection, thus leading to a lower spore count. It is also possible that the quality of the nutritional status (lipids and carbohydrates) presented by blood feeding females is different and more suitable for spore proliferation than that found in males, but this remains to be investigated.

Natural larval populations can be exposed to a range of parasite densities. The fact that neither the numbers of spores given (density of infection) nor host sex affected longevity, may in part be due to the immune response (mentioned earlier) mounted by the mosquito in response to *V. culicis*. And it is probably the initial, early, immune response that may be of more importance. So that no matter how many spores a larva ingests, the initial immune response may decimate viable spore numbers within the host to a similar level whether high or low numbers of spores have been ingested. It could also be that a ‘high spore dose’ may be much greater than the 50,000 spores per larva used here to achieve any impact on longevity or eventual spore count on death.

Infection with *V. culicis* did not affect the daily age the mosquitoes died, or the time they pupated compared to uninfected mosquitoes (Table 2, Figure 4). There was an hourly difference where infected mosquitoes died 17 to 18 hours earlier and the importance of this can only be speculated. This study was of interest because infection with the microsporidian, *V*.*culicis* hindered the subsequent development of *Plasmodium* species (Bargielowski I & Koella JC, 2009). It is possible to select for *V. culicis* strains in the laboratory that lead to earlier host death and culture such strains to achieve earlier death of the mosquito. How useful would it be for *V. culicis* or any other microsporidian to have a double effect, i.e. achieve early host death as well as impede malaria parasite development? If *V. culicis* or any other microsporidian species is highly efficient at halting the production of malaria parasites, regardless of when the larva was infected, then the timing of adult mosquito death would be inconsequential. If the suppressive effects were not the absolute halting of the secondary parasite infection, then achieving earlier host deaths would be a highly desirable microsporidian trait.

It is logical that the number of spores rose with the duration of infection (Figure 5), as the longer the time available for spore production and maturation within the host, the greater will be the total spore count, even when the more mature immune response in older hosts is factored in. This is because the mosquito’s immune reaction to *V. culicis* has been shown to be inadequate in the elimination of the parasite, hence the longer the duration of infection, the greater the spore count. However, when the same time frame was considered for spore production (i.e. comparing equal duration of infection from early and late infected larvae), the outcomes remained the same – mosquitoes infected later (older larvae) produced fewer spores (Figure 1); females produced greater numbers of spores compared to males (Figure 2); the original level of infection did not affect the spore count at death (Table 3 & Figure 3); infection did not affect the age at death of the mosquito compared to controls (Table 2 & Table 8); all of the outcomes remained the same when equal duration of infection was considered.

**Figure 2:**
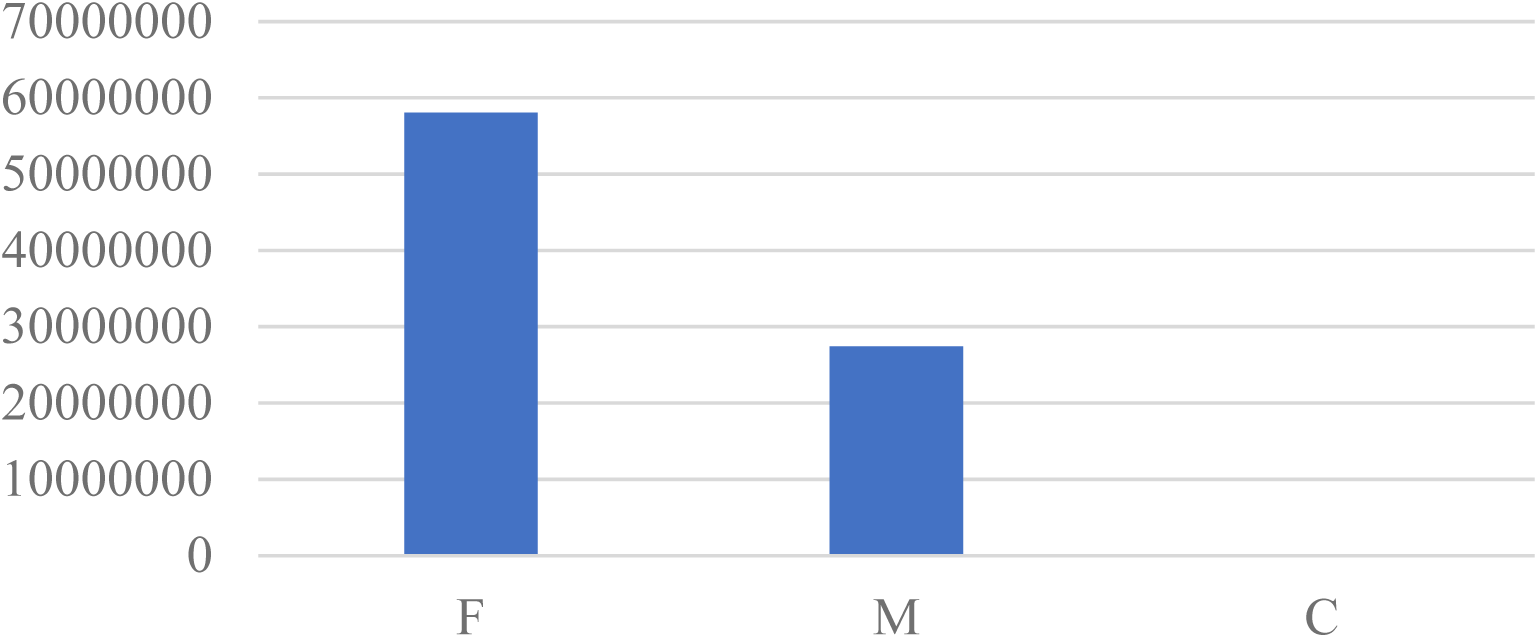
Sum of Spore.count by Sex. Females (F) produced larger numbers of spores in total than Males (M). C=Controls

**Figure 3:**
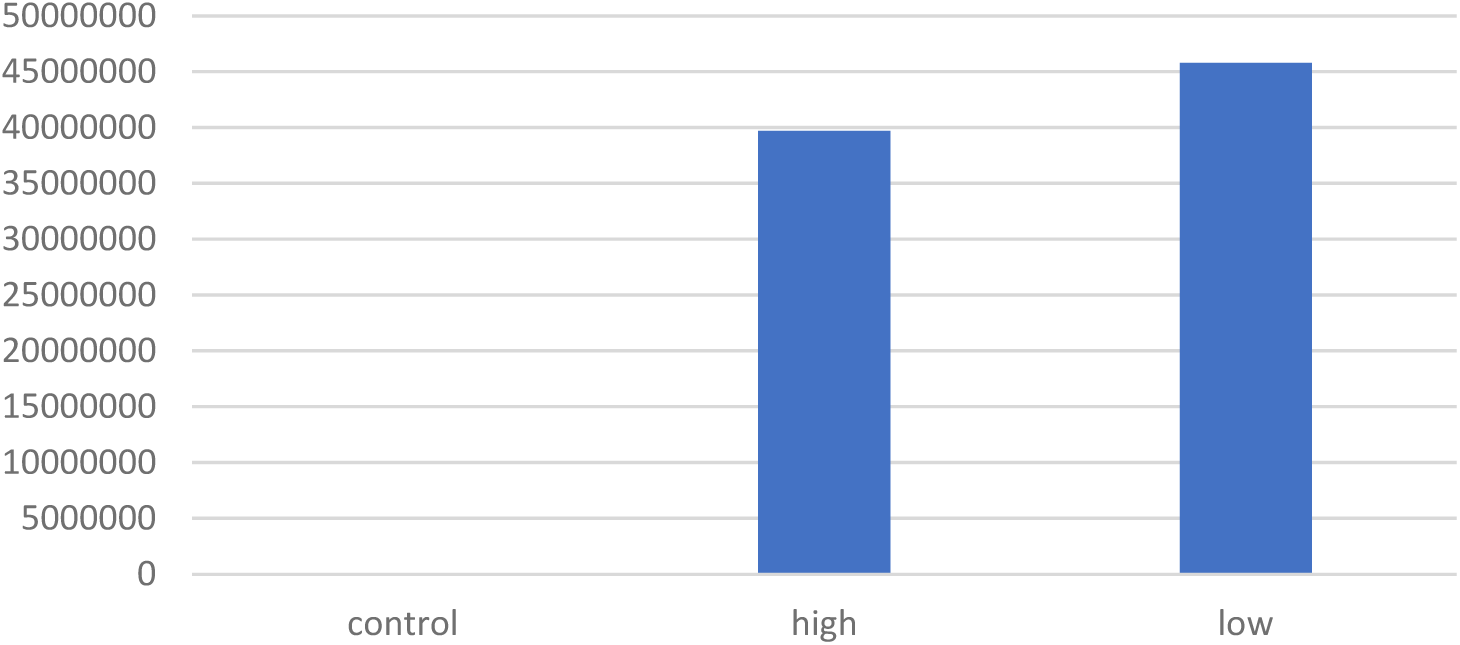
Sum of Spore.count (y axis) by Infection.level (x axis). The original level of infection in larvae did not affect the spore count in adult mosquitoes at death.

**Figure 4:**
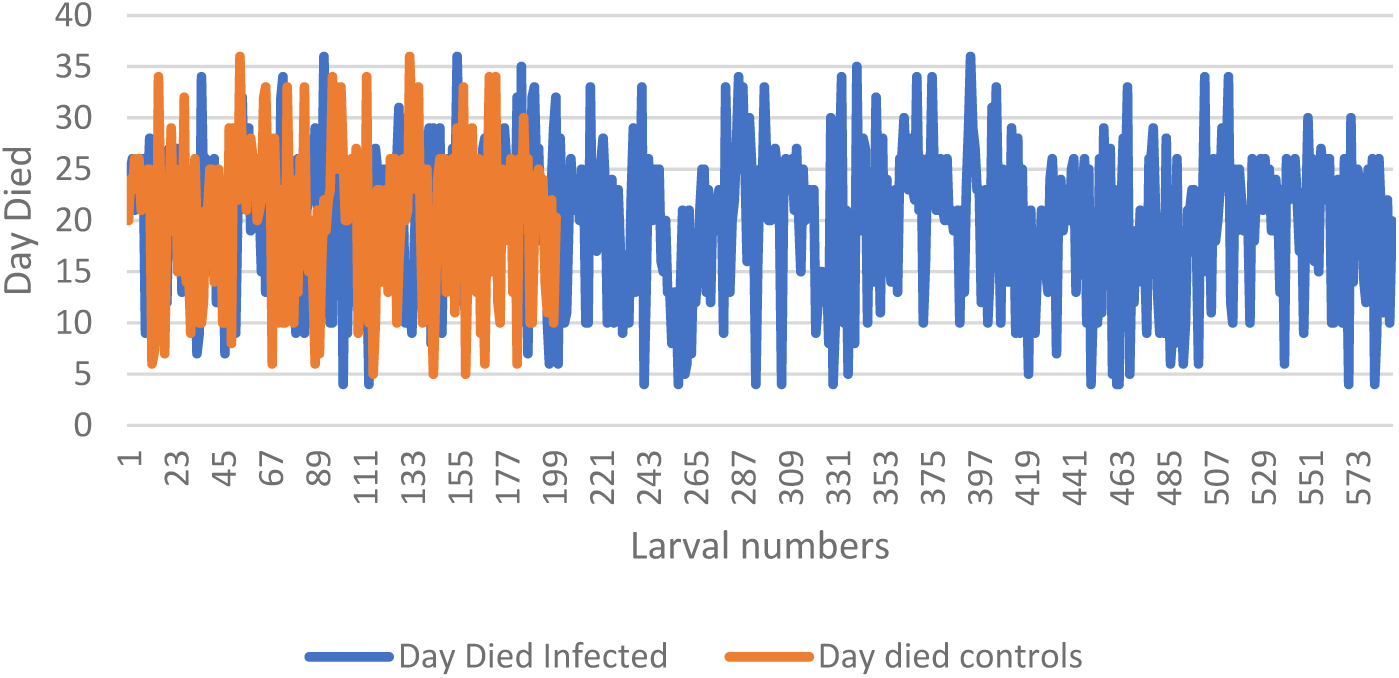
Infection does not affect the age at death of the adults.

**Figure 5:**
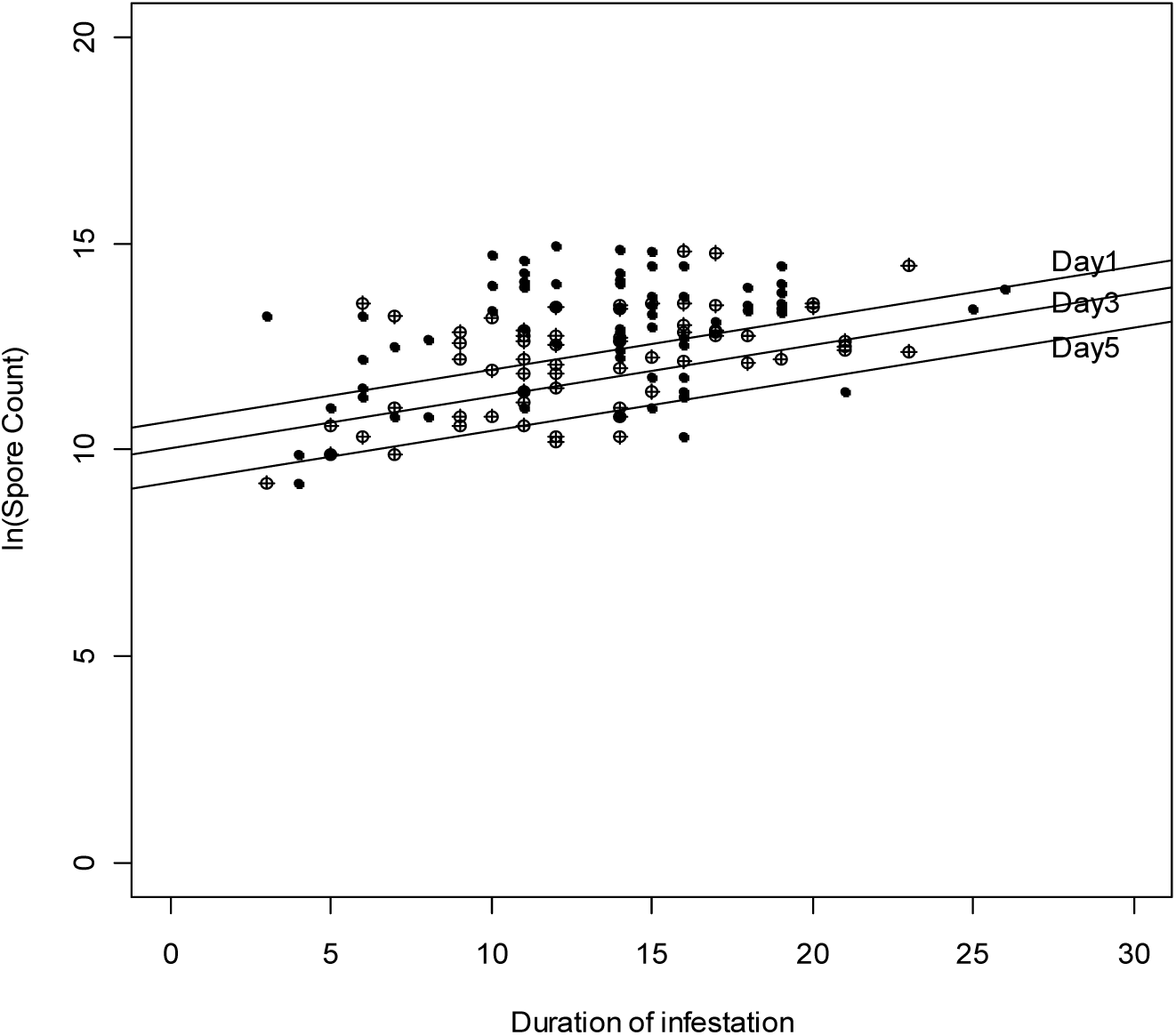
Indicating the rate of growth of *V*.*culicis* infected at different host ages, showing the initial suppression in spore production in older infected mosquitoes (lower intercepts).

Resistance to infection in older larvae has not been demonstrated in this study, lower spore production has. However, a degree of resistance to infection from older larvae is implied, as both groups received similar treatment, but older infected hosts produced fewer spores compared to younger infected larvae, indicating some resistance from older hosts. All ages and stages of the mosquito larva were susceptible to infection by *V. culicis* as spores were retrieved from all of the infected groups. Neither the younger or older infected groups could clear the infection entirely but certain host conditions as presented by younger infected larvae were more favourable for spore production. This decrease in susceptibility with larval age is in agreement with previous studies in other insect-microsporidian models (Altizer and Oberhauser 1999, Blaser M & Schmid-Hempel P 2005, Milner RJ 1973, Maddox JV et al 1990 & 2000, Vijendravarma RK et al, 2008) as well as in infections of larvae with bacterial and nematode pathogens (Umphlett CJ and Huang CS, 1973; Vyas-Patel N, 1983).

Earlier infected mosquitoes died earlier than later infected ones (although the difference here was a matter of hours) and older infected hosts produced significantly fewer spores than younger infected. This suggests that older infected larvae were less impacted than younger infected ones. In other studies, most notably *Culex pipiens* infected with *V. culicis*, mosquitoes pupated earlier and emerged as smaller adults (Agnew P et al. 1999). Here, the time of moulting did not change (Tables 4 & 5) and this may be due to species differences and the optimum food and larval rearing conditions.

It is clear that the age at which mosquitoes first encounter parasites has important implications for the development of both the parasite and the mosquito host. Host age at the time of infection is an important factor to consider in any host/parasite interaction, especially if virulence and later infections with malaria parasites is a consideration.

## Conclusions

1. The age of mosquito larvae at infection with *V. culicis* affects the spore count on death. Larvae infected later (older hosts) produced a net total of fewer spores compared to early infected, younger larvae, indicating a more robust immune response from older larvae. There was an initial delay in spore production in later infected older hosts not seen from younger infected larvae.
2. Early infected mosquitoes died slightly earlier than later infected ones. Later infected mosquitoes having between 17 to 18 hours of extra life, for every day later that they were infected.
3. The level of infection of *V. culicis*, i.e. the higher (50,000) and lower (5,000) numbers of spores supplied, did not affect the age at death of the mosquito. There were no significant interactions between infection levels (dose), sex of the mosquito and day infected. Hence the longevity of infected mosquitoes was not affected significantly by either the initial dose of *V. culicis* given or by the sex of the mosquito. This suggests a level of tolerance to the parasite by the host.
4. The level of infection with *V. culicis* (spore numbers supplied) did not affect the total spore count.
5. The age at death of the mosquitoes was not significantly affected by infection with *V. culicis*.
6. The age at pupation was not affected by the initial dose of *V. culicis* given, or by the sex of the mosquito.
7. Male mosquitoes produced fewer spores than females.
8. The number of spores from infected adults rose with increasing duration of infection.
9. These findings remained the same even when equal duration of infection was considered in both younger and older infected larvae.

## Supporting information

Statistical analysis in R (1)

Statistical analysis in R (2)

Raw data for the experiment

## Acknowledgments

This project was funded by the Daphne Jackson Trust and carried out at Imperial College; both are gratefully acknowledged. Thanks to Prof. Jacob Koella for reading and commenting on the manuscript and to Dr Catherine M Collins for valuable help with the statistical analysis.

## References

Agnew P, Bedhomme S, Haussy C, Michalakis Y, 1999. Age and size at maturity of the mosquito Culex pipiens infected by the microsporidian parasite Vavraia culicis. Proceedings of the Royal Society of London Series B-Biological Sciences. 1999;266:947–952. https://doi.org/10.1098/rspb.1999.0728; https://royalsocietypublishing.org/doi/abs/10.1098/rspb.1999.0728.

Altizer, SM and Oberhauser K.S. 1999. Effects of the protozoan parasite Ophryocystis elektroscirrha on the fitness of monarch butterflies (Danaus plexippus). Journal of Invertebrate Pathology 74 (1): 76–88. https://www.sciencedirect.com/science/article/abs/pii/S002220119994853X?via%3Dihub

Andreadis TG, 2007. Microsporidian parasites of mosquitoes. Journal of the American Mosquito Control Association 23(2 Suppl):3–29 DOI: 10.2987/8756-971X(2007)23[3:MPOM]2.0.CO;2 https://www.researchgate.net/publication/5986217_Microsporidian_parasites_of_mosquitoes

Bano L 1958. Partial Inhibitory Effect of Plistophora culicis on the Sporogonic Cycle of of Plasmodium cynomolgi in Anopheles Stephensi Nature volume181, page430 (1958). https://www.nature.com/articles/181430a0

Bargielowski I & Koella JC, 2009. A Possible Mechanism for the Suppression of Plasmodium berghei Development in the Mosquito Anopheles gambiae by the Microsporidian Vavraia culici. PLoS ONE 4(3): e4676. doi:10.1371/journal.pone.0004676. https://journals.plos.org/plosone/article?id=10.1371/journal.pone.0004676

Bedhomme S, Agnew P, Sidobre, S, Michalakis Y 2004. Virulence reaction norms across a food gradient. Proc. Royal Soc. Biological Sciences. 271 (1540): 739–744. https://royalsocietypublishing.org/doi/10.1098/rspb.2003.2657

Biron DG, Agnew P, Marche L, Renault L, Sidobre C, and Michalakis Y, 2005. Proteome of Aedes aegypti larvae in response to infection by the intracellular parasite Vavraia culicis. International Journal for Parasitology 35: 1385–1397. https://www.sciencedirect.com/science/article/abs/pii/S0020751905002067?via%3Dihub

Blaser M and Schmid-Hempel P, 2005. Determinants of virulence for the parasite Nosema whitei in its host Tribolium castaneum. Journal of Invertebrate Pathology 89 (3): 251–257. https://doi.org/10.1016/j.jip.2005.04.004https://www.sciencedirect.com/science/article/abs/pii/S0022201105000637?via%3Dihub

Brown LD, Shapiro LLM, Thompson GA, Estévez-Lao TY, and Hillyer JF, 2019. Transstadial immune activation in a mosquito: Adults that emerge from infected larvae have stronger antibacterial activity in their hemocoel yet increased susceptibility to malaria infection. Ecol Evol. 2019 May; 9(10): 6082–6095. doi: 10.1002/ece3.5192. PMCID: PMC6540708 PMID: 31161020. https://www.ncbi.nlm.nih.gov/pmc/articles/PMC6540708/

Cheney SA, Lafranchi-Tristem NJ, Canning EU, 2005. Phylogenetic relationship of Pleisotophora-like microsporidia based on small subunit ribosomal DNA sequences and implications for the source of Trachpleistophora hominis infections. J Euk Microbiol 47:280–287. https://doi.org/10.1111/j.1550-7408.2000.tb00048.x. https://onlinelibrary.wiley.com/doi/abs/10.1111/j.1550-7408.2000.tb00048.x.

Futuyma DJ, 2001. Co-Evolution, Evolution of Virulence and Avirulence, pp 407 to 412. In Encyclopedia of Genetics, Editors Brenner S & Miller JH. 2,800 pp. ISBN 978-0-12-227080-2. Elsevier Science. https://www.sciencedirect.com/science/article/pii/B0122270800002391

Habtewold T, Duchateau L and Christophides GK, 2016. Flow Cytometry analysis of the microbiota associated with the midguts of vector mosquitoes. Parasites and Vectors 2016, Vol: 9, ISSN: 1756-3305. http://dx.doi.org/10.1186/s13071-016-1438-0. https://parasitesandvectors.biomedcentral.com/articles/10.1186/s13071-016-1438-0

Kelly JF, Anthony DW, Dillard CR, 1981. A laboratory evaluation of the microsporidian Vavraia culicis as an agent for mosquito control. Journal of Invertebrate Pathology. 1981;37:117–122. https://doi.org/10.1016/0022-2011(81)90064-1; https://www.sciencedirect.com/science/article/abs/pii/0022201181900641

League GP, Estévez-Lao TY, Yan Y, Garcia-Lopez VA and Hillyer JF, 2017. Anopheles gambiae larvae mount stronger immune responses against bacterial infection than adults: evidence of adaptive decoupling in mosquitoes. Parasites & Vectors (2017) 10:367. DOI 10.1186/s13071-017-2302-6. https://d-nb.info/1141733439/34 https://www.ncbi.nlm.nih.gov/pmc/articles/PMC5539753/

Li X, Palmer R, Trout JM, and Fayer R 2003. Infectivity of Micropsoridia spores stored in water at Environmental Temperatures. J. Parasitol., 89(1), 2003,pp. 185-188. American Society of Parasitologists 2003. https://naldc.nal.usda.gov/download/21417/PDF?ev=pub_ext_prw_xdl

Lorenz LM & Koella JC, 2011. The microsporidian parasite Vavraia culicis as a potential late life–acting control agent of malaria. Evol Appl. 2011 Nov; 4(6): 783–790. doi: 10.1111/j.1752-4571.2011.00199.x. https://www.ncbi.nlm.nih.gov/pmc/articles/PMC3352544/.

Lounibos LP, Nishimura N, Conn J, Lourenco-de-Oliveira R, 1995. Life History Correlates of Adult Size in the Malaria Vector Anopheles darlingi. Memorias do Instituto Oswaldo Cruz, 90 (6). https://www.scielo.br/pdf/mioc/v90n6/vol90(f6)_099-104.pdf

Maddox JV, Brooks WM and Solter LF (2000). Bioassays of microsporidia: in bioassays of entomopathogenic microbes and nematodes. Pages 197–228. https://www.cabi.org/cabebooks/ebook/20001108781

Michalakis Y, Bedhomme S, Biron DG, Rivero A, Sidobre C, Agnew P, 2008. Virulence and Resistance in a Mosquito-Microsporidian Interaction. Journal Compilation Copyright 2008, Blackwell Publishing Ltd. 1 (2008) 49–56. doi: 10.1111/j.1752-4571.2007.00004.x https://www.ncbi.nlm.nih.gov/pmc/articles/PMC3352405/

Milner R J, 1973. Nosema whitei, a microsporidan pathogen of some species of Tribolium. IV. The effect of temperature, humidity and larval age on pathogenicity for T. castaneum. Entomophaga 18 (3): 305–315. http://citeseerx.ist.psu.edu/viewdoc/download?doi=10.1.1.547.8537&rep=rep1&type=pdf

Onstad, D.W and Maddox JV. 1990. Simulation-model of Tribolium confusum and its Pathogen, Nosema whitei. Ecological Modelling 51 (1-2): 143–160. https://www.sciencedirect.com/science/article/abs/pii/030438009090062L

Reynolds DG, 1970. Laboratory studies of the microsporidian Plistophora culicis (Weiser) infecting Culex pipiens fatigans Wied. Bull Entomol Res. 1970 Dec; 60(2):339–49. https://www.ncbi.nlm.nih.gov/pubmed/22894851/

Rivero A, Agnew P, Bedhomme S, Sidobre C, and Michalakis Y, 2007. Resource depletion in Aedes aegypti mosquitoes infected by the microsporidia Vavraia culicis. Parasitology, 134:1355–1362. DOI: https://doi.org/10.1017/S0031182007002703 https://www.researchgate.net/publication/6204558_Resource_depletion_in_Aedes_aegypti_mosquitoes_infected_by_the_microsporidia_Vavraia_culicis

Roberts D, 2014. Mosquito Larvae Change Their Feeding Behavior in Response to Kairomones From Some Predators. Journal of Medical Entomology, Volume 51, Issue 2, 1 March 2014, Pages 368–374, https://doi.org/10.1603/ME13129; https://academic.oup.com/jme/article/51/2/368/882256.

Thornhill R, Alcock J, 1983. The Evolution of Insect Mating Systems. Harvard University Press, Cambridge, Massachusetts, ix+547pp. https://onlinelibrary.wiley.com/doi/abs/10.1111/j.1558-5646.1984.tb00343.x

Umphlett CJ and Huang CS, 1972. Experimental infection of mosquito larvae by a species of the aquatic fungus Lagenidium JIP 20, 3 1972, Pp 326–331. https://doi.org/10.1016/0022-2011(72)90164-4. https://www.sciencedirect.com/science/article/abs/pii/0022201172901644.

Vávra J and Becnel JJ, 2007. Vavraia culicis (Weiser, 1947 & 1977) revisited: cytological characterisation of a Vavraia culicis-like microsporidium isolated from mosquitoes in Florida and the establishment of Vavraia culicis floridensis subsp. n. Folia Parasitologica 54 (4) 259–271, 2007. DOI: 10.14411/fp.2007.034. https://folia.paru.cas.cz/artkey/fol-200704-0002_Vavraia_culicis_Weiser_1947_Weiser_1977_revisited_cytological_characterisation_of_a_Vavraia_culicis-like_m.php

Vijendravarma RK, Godfray CH, & Kraaijeveld AR, 2008. Infection of Drosophila melanogaster by Tubulinosema kingi: Stage-specific susceptibility and within-host proliferation. JIP 99 239–241. https://doi.org/10.1016/j.jip.2008.02.014 https://www.sciencedirect.com/science/article/abs/pii/S0022201108000566?via%3Dihub

Vyas-Patel N, 1988. Parasitism of Kenyan Mosquito Larvae (Diptera: Culicidae) by Romanomermis culicivorax (Nematoda: Mermithidae). J Nematol. 1988 Jan; 20(1): 96–101. PMCID: PMC2618791, PMID: 19290190 https://www.ncbi.nlm.nih.gov/pmc/articles/PMC2618791/

Vyas-Patel N, 1983. Effect of Aedes Aegypti (Diptera:Culicidae) Age on Sex Ratios in Romanomermis Culicivorax (Nematoda:Mermithidae). J Nematol 1983 Oct;15(4):594-7. J Nematol. 1983 Oct; 15(4): 594–597. PMID: 19295853 PMCID: PMC2618322. https://www.ncbi.nlm.nih.gov/pmc/articles/PMC2618322/.

Vyas-Patel N, 2019. Monitoring transgenic mosquitoes using wing measurements and I3S Classic. BioRxiv May 23rd, 2019. doi: https://doi.org/10.1101/645028. https://www.biorxiv.org/content/10.1101/645028v1#disqus_thread.

Weiser J, 1947. A key for the determination of microsporidia. Práce Mor. Přír. Spol. 18: 1–64. (In Czech.). https://naldc.nal.usda.gov/download/55882/PDF

Weiser J, 1977. Contribution to the classification of microsporidia. Věstn. Čs. Spol. Zool. 41: 308–320.

Weiser J, 1969. Immunity of insects to protozoa. in Immunity to Parasitic Animals. 1: 129–147.

