## Supplementary material for "*Anopheles coluzzii* infection by the microsporidian, *Vavraia culicis*: the effect of host age": Statistical analysis in R (1)

```
---
R version 2.10.1 (2009-12-14)
Copyright (C) 2009 The R Foundation for Statistical Computing
ISBN 3-900051-07-0

R is free software and comes with ABSOLUTELY NO WARRANTY.
You are welcome to redistribute it under certain conditions.
Type 'license()' or 'licence()' for distribution details.

  Natural language support but running in an English locale

R is a collaborative project with many contributors.
Type 'contributors()' for more information and
'citation()' on how to cite R or R packages in publications.

Type 'demo()' for some demos, 'help()' for on-line help, or
'help.start()' for an HTML browser interface to help.
Type 'q()' to quit R.

[Previously saved workspace restored]

> dur<-read.table("D://Dura.txt",Header=T)
Error in read.table("D://Dura.txt", Header = T) :
  unused argument(s) (Header = T)
> dur<-read.table("D://Dura.txt",header=T)
> names(dur)
[1] "Infection.level" "Day.Infected"    "Spore.count"     "Duration"
> attach(dur)
> mod1<-glm(Spore.count~Day.Infected*Duration*Infection.level,quasipoisson)
Error: NA/NaN/Inf in foreign function call (arg 1)
> rm(dur)
> dur<-read.table("D://Dura.txt",Header=T)
Error in read.table("D://Dura.txt", Header = T) :
  unused argument(s) (Header = T)
> dur<-read.table("D://Dura.txt",header=T)
> attach(dur)

        The following object(s) are masked from dur ( position 3 ) :

         Day.Infected Duration Infection.level Spore.count

> mod1<-glm(Spore.count~Day.Infected*Duration*Infection.level,quasipoisson)
Error: NA/NaN/Inf in foreign function call (arg 1)
> hist(Spore.count)
> levels(Day.Infected)
NULL
> levels(Infection.level)
[1] "high" "low"
> mod1<-glm(Spore.count~Day.Infected*Duration+Infection.level,quasipoisson)
Error: NA/NaN/Inf in foreign function call (arg 1)
In addition: Warning message:
step size truncated due to divergence
> mod1<-glm(Spore.count~Day.Infected+Duration+Infection.level,quasipoisson)
```

```
> summary(mod1)

Call:
glm(formula = Spore.count ~ Day.Infected + Duration + Infection.level,
    family = quasipoisson)

Deviance Residuals:
     Min        1Q    Median        3Q       Max
-1216.169  -384.525   -60.366     1.110  2586.743

Coefficients:
                  Estimate Std. Error   t value Pr(>|t|)
(Intercept)       -6.483e+00  2.517e+04 -2.58e-04    1.000
Day.Infected      -3.846e-01  5.773e-02    -6.662 1.53e-10 ***
Duration1         -3.073e-01  3.060e+04 -1.00e-05    1.000
Duration10         1.837e+01  2.517e+04     0.001    0.999
Duration11         1.581e+01  2.517e+04     0.001    0.999
Duration12         1.694e+01  2.517e+04     0.001    0.999
Duration13         1.946e+01  2.517e+04     0.001    0.999
Duration14         1.865e+01  2.517e+04     0.001    0.999
Duration15         1.914e+01  2.517e+04     0.001    0.999
Duration16         1.961e+01  2.517e+04     0.001    0.999
Duration17         2.055e+01  2.517e+04     0.001    0.999
Duration18         2.048e+01  2.517e+04     0.001    0.999
Duration19         2.027e+01  2.517e+04     0.001    0.999
Duration2          1.036e-01  3.560e+04  2.91e-06    1.000
Duration20         2.022e+01  2.517e+04     0.001    0.999
Duration21         2.028e+01  2.517e+04     0.001    0.999
Duration22         2.022e+01  2.517e+04     0.001    0.999
Duration23         2.085e+01  2.517e+04     0.001    0.999
Duration24         2.049e+01  2.517e+04     0.001    0.999
Duration25         2.060e+01  2.517e+04     0.001    0.999
Duration26         2.124e+01  2.517e+04     0.001    0.999
Duration27         1.969e+01  2.517e+04     0.001    0.999
Duration28         1.821e+01  2.517e+04     0.001    0.999
Duration29         2.151e+01  2.517e+04     0.001    0.999
Duration3         -1.148e+00  2.593e+04 -4.43e-05    1.000
Duration31         2.018e+01  2.517e+04     0.001    0.999
Duration33         2.068e+01  2.517e+04     0.001    0.999
Duration4          4.201e-02  2.669e+04  1.57e-06    1.000
Duration5         -4.375e-01  2.778e+04 -1.57e-05    1.000
Duration6         -5.868e-01  2.752e+04 -2.13e-05    1.000
Duration7         -6.169e-01  2.715e+04 -2.27e-05    1.000
Duration8         -8.887e-01  2.708e+04 -3.28e-05    1.000
Duration9         -8.021e-01  2.651e+04 -3.03e-05    1.000
Durationna         7.692e-01  3.560e+04  2.16e-05    1.000
Infection.levellow  1.036e-01  1.490e-01     0.695    0.488
---
Signif. codes:  0 ‘***’ 0.001 ‘**’ 0.01 ‘*’ 0.05 ‘.’ 0.1 ‘ ’ 1

(Dispersion parameter for quasipoisson family taken to be 426999.4)

    Null deviance: 189011396  on 302  degrees of freedom
Residual deviance:  88621301  on 268  degrees of freedom
```

```
  (97 observations deleted due to missingness)
AIC: NA

Number of Fisher Scoring iterations: 6

> rm(dur)
> dur<-read.table("D://Dura.txt",header=T)
> attach(dur)

        The following object(s) are masked from dur ( position 3 ) :

         Day.Infected Duration Infection.level Spore.count


        The following object(s) are masked from dur ( position 4 ) :

         Day.Infected Duration Infection.level Spore.count

> mod1<-glm(Spore.count~Day.Infected+Duration+Infection.level,quasipoisson)
> summary(mod1)

Call:
glm(formula = Spore.count ~ Day.Infected + Duration + Infection.level,
    family = quasipoisson)

Deviance Residuals:
     Min        1Q    Median        3Q       Max
-1393.43   -452.57   -298.35     23.59   2449.60

Coefficients:
                   Estimate Std. Error t value Pr(>|t|)
(Intercept)        11.06549    0.34975  31.638  < 2e-16 ***
Day.Infected       -0.33046    0.05592  -5.909 9.39e-09 ***
Duration            0.12384    0.01453   8.524 7.85e-16 ***
Infection.levellow  0.03805    0.15968   0.238    0.812
---
Signif. codes:  0 '***' 0.001 '**' 0.01 '*' 0.05 '.' 0.1 ' ' 1

(Dispersion parameter for quasipoisson family taken to be 527812.5)

    Null deviance: 188452473  on 301  degrees of freedom
Residual deviance: 114003175  on 298  degrees of freedom
AIC: NA

Number of Fisher Scoring iterations: 6

> mod2<-glm(Spore.count~Day.Infected+Duration,quasipoisson)
> summary(mod2)

Call:
glm(formula = Spore.count ~ Day.Infected + Duration, family = quasipoisson)

Deviance Residuals:
     Min        1Q    Median        3Q       Max
```

```
-1413.60   -452.50   -309.00     29.06   2466.23  

Coefficients:
             Estimate Std. Error t value Pr(>|t|)    
(Intercept)  11.07790    0.34647  31.974  < 2e-16 ***
Day.Infected -0.33075    0.05595  -5.912 9.23e-09 ***
Duration      0.12425    0.01447   8.586 5.03e-16 ***
---
Signif. codes:  0 ‘***’ 0.001 ‘**’ 0.01 ‘*’ 0.05 ‘.’ 0.1 ‘ ’ 1 

(Dispersion parameter for quasipoisson family taken to be 528537.5)

    Null deviance: 188452473  on 301  degrees of freedom
Residual deviance: 114033169  on 299  degrees of freedom
AIC: NA

Number of Fisher Scoring iterations: 6

> DayF<-as.factor(Day,Infected)
Error in as.factor(Day, Infected) : unused argument(s) (Infected)
> DayF<-as.factor(Day.Infected)
> mad2<-glm(Spore.count~DayF*Duration,quasipoisson)
> summary(mad2)

Call:
glm(formula = Spore.count ~ DayF * Duration, family = quasipoisson)

Deviance Residuals: 
     Min        1Q    Median        3Q       Max  
-1303.50   -452.85   -235.28     24.61   2424.54  

Coefficients:
               Estimate Std. Error t value Pr(>|t|)    
(Intercept)    11.23360    0.36316  30.933  < 2e-16 ***
DayF3          -1.99318    0.72432  -2.752  0.00629 ** 
DayF5          -2.98275    1.06801  -2.793  0.00557 ** 
DayF7          -2.24861    3.62830  -0.620  0.53591    
Duration        0.10146    0.01696   5.980 6.45e-09 ***
DayF3:Duration  0.06321    0.03293   1.919  0.05589 .  
DayF5:Duration  0.07301    0.05046   1.447  0.14901    
DayF7:Duration  0.11006    0.20294   0.542  0.58801    
---
Signif. codes:  0 ‘***’ 0.001 ‘**’ 0.01 ‘*’ 0.05 ‘.’ 0.1 ‘ ’ 1 

(Dispersion parameter for quasipoisson family taken to be 477229.4)

    Null deviance: 188452473  on 301  degrees of freedom
Residual deviance: 109003146  on 294  degrees of freedom
AIC: NA

Number of Fisher Scoring iterations: 7

> mad3<-glm(Spore.count~DayF+Duration+Infection.level,quasipoisson)
> summary(mad3)
```

```
Call:
glm(formula = Spore.count ~ DayF + Duration + Infection.level,
    family = quasipoisson)

Deviance Residuals:
     Min        1Q    Median        3Q       Max
-1413.78   -435.90   -284.04     24.77   2448.77

Coefficients:
                   Estimate Std. Error t value Pr(>|t|)
(Intercept)        10.73619    0.31742  33.824  < 2e-16 ***
DayF3              -0.65983    0.17216  -3.833 0.000155 ***
DayF5              -1.50242    0.25885  -5.804 1.66e-08 ***
DayF7              -0.27925    0.63934  -0.437 0.662589
Duration            0.12466    0.01404   8.880  < 2e-16 ***
Infection.levellow  0.01863    0.15392   0.121 0.903748
---
Signif. codes:  0 ‘***’ 0.001 ‘**’ 0.01 ‘*’ 0.05 ‘.’ 0.1 ‘ ’ 1

(Dispersion parameter for quasipoisson family taken to be 487394)

    Null deviance: 188452473  on 301  degrees of freedom
Residual deviance: 111515775  on 296  degrees of freedom
AIC: NA

Number of Fisher Scoring iterations: 6

> mad4<-glm(Spore.count~DayF+Duration,quasipoisson)
> summary(mad4)

Call:
glm(formula = Spore.count ~ DayF + Duration, family = quasipoisson)

Deviance Residuals:
     Min        1Q    Median        3Q       Max
-1423.73   -434.55   -285.06     30.05   2456.88

Coefficients:
            Estimate Std. Error t value Pr(>|t|)
(Intercept) 10.74193    0.31375  34.237  < 2e-16 ***
DayF3       -0.66002    0.17198  -3.838 0.000152 ***
DayF5       -1.50327    0.25849  -5.816 1.56e-08 ***
DayF7       -0.27319    0.63677  -0.429 0.668217
Duration     0.12487    0.01394   8.960  < 2e-16 ***
---
Signif. codes:  0 ‘***’ 0.001 ‘**’ 0.01 ‘*’ 0.05 ‘.’ 0.1 ‘ ’ 1

(Dispersion parameter for quasipoisson family taken to be 486432.7)

    Null deviance: 188452473  on 301  degrees of freedom
Residual deviance: 111522917  on 297  degrees of freedom
AIC: NA
```

```
Number of Fisher Scoring iterations: 6

> tapply(Spore.count,DayF,length)
  1   3   5   7 
 99 100  98   5 
> plot(Spore.count~Duration)
> plot(log(Spore.count)~Duration)
> abline(10.73,0.13)
> abline((10.73-0.66),0.13)
> abline((10.73-1.503),0.13)
> plot(Spore.count~Duration)
> plot(log(Spore.count)~Duration)
> abline(10.73,0.13)
> abline((10.73-0.66),0.13)
> abline((10.73-1.503),0.13)
> summary(Duration)
   Min. 1st Qu.  Median    Mean 3rd Qu.    Max. 
   0.00   11.00   17.00   15.94   21.00   33.00 
> summary(mod2)

Call:
glm(formula = Spore.count ~ Day.Infected + Duration, family = quasipoisson)

Deviance Residuals: 
     Min        1Q    Median        3Q       Max  
-1413.60   -452.50   -309.00     29.06   2466.23  

Coefficients:
             Estimate Std. Error t value Pr(>|t|)    
(Intercept)  11.07790    0.34647  31.974  < 2e-16 ***
Day.Infected -0.33075    0.05595  -5.912 9.23e-09 ***
Duration      0.12425    0.01447   8.586 5.03e-16 ***
---
Signif. codes:  0 ‘***’ 0.001 ‘**’ 0.01 ‘*’ 0.05 ‘.’ 0.1 ‘ ’ 1

(Dispersion parameter for quasipoisson family taken to be 528537.5)

    Null deviance: 188452473  on 301  degrees of freedom
Residual deviance: 114033169  on 299  degrees of freedom
AIC: NA

Number of Fisher Scoring iterations: 6

> summary(mod1)

Call:
glm(formula = Spore.count ~ Day.Infected + Duration + Infection.level, 
    family = quasipoisson)

Deviance Residuals: 
     Min        1Q    Median        3Q       Max  
-1393.43   -452.57   -298.35     23.59   2449.60  

Coefficients:
```

```
                 Estimate Std. Error t value Pr(>|t|)
(Intercept)       11.06549    0.34975  31.638  < 2e-16 ***
Day.Infected      -0.33046    0.05592  -5.909 9.39e-09 ***
Duration           0.12384    0.01453   8.524 7.85e-16 ***
Infection.levellow  0.03805    0.15968   0.238    0.812
---
Signif. codes:  0 ‘***’ 0.001 ‘**’ 0.01 ‘*’ 0.05 ‘.’ 0.1 ‘ ’ 1

(Dispersion parameter for quasipoisson family taken to be 527812.5)

    Null deviance: 188452473  on 301  degrees of freedom
Residual deviance: 114003175  on 298  degrees of freedom
AIC: NA

Number of Fisher Scoring iterations: 6

> plot(Spore.count~Duration)
> rm(dur)
> nan<-read.table("D://Nayna.txt",header=T)
> attach(nan)

        The following object(s) are masked from dur ( position 3 ) :

         Day.Infected Duration Infection.level Spore.count


        The following object(s) are masked from dur ( position 4 ) :

         Day.Infected Duration Infection.level Spore.count


        The following object(s) are masked from dur ( position 5 ) :

         Day.Infected Duration Infection.level Spore.count

> names(nan)
[1] "Plate"           "Infection.level" "Day.Infected"
[4] "Pupated"         "Died"            "Sex"
[7] "Wing.length"     "Spore.count"     "Duration"
> boxplot(Pupated~Infection.level)
> tapply(Pupated,Infection.level,mean)
control    high     low
     NA      NA      NA
> pup1<-aov(Pupated~Infection.level)
> summary(pup1)
                 Df Sum Sq Mean Sq F value  Pr(>F)
Infection.level   2   1.44 0.72002  2.6371 0.07245 .
Residuals       564 153.99 0.27304
---
Signif. codes:  0 ‘***’ 0.001 ‘**’ 0.01 ‘*’ 0.05 ‘.’ 0.1 ‘ ’ 1
33 observations deleted due to missingness
> pup2<-aov(Pupated~Infection.level+Day.Infected)
> summary(pup2)
                 Df Sum Sq Mean Sq F value  Pr(>F)
```

```
Infection.level   1  0.421 0.42064  1.7101 0.19177
Day.Infected      1  1.554 1.55435  6.3194 0.01236 *
Residuals       375 92.237 0.24596
---
Signif. codes:  0 ‘***’ 0.001 ‘**’ 0.01 ‘*’ 0.05 ‘.’ 0.1 ‘ ’ 1
222 observations deleted due to missingness
> pup3<-aov(Pupated~Infection.level*Day.Infected)
>
> summary(pup2-3)
Error in pup2 - 3 : non-numeric argument to binary operator
> summary(pup3)
                             Df Sum Sq Mean Sq F value  Pr(>F)
Infection.level               1  0.421 0.42064  1.7143 0.19123
Day.Infected                  1  1.554 1.55435  6.3349 0.01226 *
Infection.level:Day.Infected   1  0.471 0.47132  1.9209 0.16658
Residuals                   374 91.765 0.24536
---
Signif. codes:  0 ‘***’ 0.001 ‘**’ 0.01 ‘*’ 0.05 ‘.’ 0.1 ‘ ’ 1
222 observations deleted due to missingness
> pup4<-aov(Pupated~Day.Infected)
> summary(pup4)
              Df Sum Sq Mean Sq F value  Pr(>F)
Day.Infected   1  1.602  1.6017  6.5028 0.01117 *
Residuals    376 92.610  0.2463
---
Signif. codes:  0 ‘***’ 0.001 ‘**’ 0.01 ‘*’ 0.05 ‘.’ 0.1 ‘ ’ 1
222 observations deleted due to missingness
> boxplot(Pupated~Day.Infected)
> plot(Pupated~Day.Infected)
> pup5<-lm(Pupated~Day.Infected)
> summary(pup5)

Call:
lm(formula = Pupated ~ Day.Infected)

Residuals:
    Min      1Q  Median      3Q     Max
-1.2436 -0.2436 -0.1850 -0.1263  1.8150

Coefficients:
             Estimate Std. Error t value Pr(>|t|)
(Intercept)   7.09701    0.05351  132.63   <2e-16 ***
Day.Infected  0.02932    0.01150    2.55   0.0112 *
---
Signif. codes:  0 ‘***’ 0.001 ‘**’ 0.01 ‘*’ 0.05 ‘.’ 0.1 ‘ ’ 1

Residual standard error: 0.4963 on 376 degrees of freedom
  (222 observations deleted due to missingness)
Multiple R-squared: 0.017,      Adjusted R-squared: 0.01439
F-statistic: 6.503 on 1 and 376 DF,  p-value: 0.01117

> pup6<-aov(Pupated~Infection.level+Day.Infected+Sex)
> summary(pup6)
                 Df Sum Sq Mean Sq F value   Pr(>F)
```

```
Infection.level   1  0.459 0.45868  1.8225 0.177886
Day.Infected      1  1.908 1.90845  7.5829 0.006198 **
Sex               1  0.511 0.51090  2.0300 0.155112
Residuals       352 88.591 0.25168
---
Signif. codes:  0 ‘***’ 0.001 ‘**’ 0.01 ‘*’ 0.05 ‘.’ 0.1 ‘ ’ 1
244 observations deleted due to missingness
> abline(7.1,0.03)
> tapply(Pupated,Day.Infected,mean)
 1  3  5  7
NA NA NA NA
> tapply(Pupated,Day.Infected,sum)
 1  3  5  7
NA NA NA NA
> names(nan)
[1] "Plate"           "Infection.level" "Day.Infected"
[4] "Pupated"         "Died"            "Sex"
[7] "Wing.length"     "Spore.count"     "Duration"
> hist(Died)
> hist(Died,breaks=20)
> ded1<-aov(Died~Infection.level*Day.Infected*Sex)
> summary(ded1)
                                 Df  Sum Sq Mean Sq F value
Infection.level                   1    14.2   14.18  0.4810
Day.Infected                      1   625.5  625.50 21.2169
Sex                               1     2.0    1.98  0.0672
Infection.level:Day.Infected      1     1.6    1.59  0.0541
Infection.level:Sex               1     8.7    8.65  0.2935
Day.Infected:Sex                  1     0.7    0.70  0.0237
Infection.level:Day.Infected:Sex   1    46.9   46.95  1.5925
Residuals                       349 10289.0   29.48
                                   Pr(>F)
Infection.level                    0.4884
Day.Infected                    5.756e-06 ***
Sex                                0.7956
Infection.level:Day.Infected       0.8162
Infection.level:Sex                0.5883
Day.Infected:Sex                   0.8778
Infection.level:Day.Infected:Sex    0.2078
Residuals
---
Signif. codes:  0 ‘***’ 0.001 ‘**’ 0.01 ‘*’ 0.05 ‘.’ 0.1 ‘ ’ 1
243 observations deleted due to missingness
> summary(ded1)
                                 Df  Sum Sq Mean Sq F value    Pr(>F)
Infection.level                   1    14.2   14.18  0.4810    0.4884
Day.Infected                      1   625.5  625.50 21.2169 5.756e-06 ***
Sex                               1     2.0    1.98  0.0672    0.7956
Infection.level:Day.Infected      1     1.6    1.59  0.0541    0.8162
Infection.level:Sex               1     8.7    8.65  0.2935    0.5883
Day.Infected:Sex                  1     0.7    0.70  0.0237    0.8778
Infection.level:Day.Infected:Sex   1    46.9   46.95  1.5925    0.2078
Residuals                       349 10289.0   29.48
---
```

```
Signif. codes:  0 ‘***’ 0.001 ‘**’ 0.01 ‘*’ 0.05 ‘.’ 0.1 ‘ ’ 1
243 observations deleted due to missingness
> ded2<-updateded1~.-Infection.level:Day.Infected:Sex)
Error: unexpected ')' in "ded2<-updateded1~.-Infection.level:Day.Infected:Sex)"
> ded2<-update(ded1~.-Infection.level:Day.Infected:Sex)
Error in inherits(object, "formula") : element 1 is empty;
   the part of the args list of '.Internal' being evaluated was:
   (x, what, which)
> ded2<-update(ded1,~.-Infection.level:Day.Infected:Sex)
> summary(ded2)
                            Df  Sum Sq Mean Sq F value    Pr(>F)
Infection.level              1    14.2   14.18  0.4802    0.4888
Day.Infected                 1   625.5  625.50 21.1810 5.853e-06 ***
Sex                          1     2.0    1.98  0.0671    0.7957
Infection.level:Day.Infected   1     1.6    1.59  0.0540    0.8164
Infection.level:Sex          1     8.7    8.65  0.2930    0.5886
Day.Infected:Sex             1     0.7    0.70  0.0236    0.8779
Residuals                  350 10336.0   29.53
---
Signif. codes:  0 ‘***’ 0.001 ‘**’ 0.01 ‘*’ 0.05 ‘.’ 0.1 ‘ ’ 1
243 observations deleted due to missingness
> ded3<-update(ded2,~.-Day.Infected:Sex)
> summary(ded3)
                            Df  Sum Sq Mean Sq F value   Pr(>F)
Infection.level              1    14.2   14.18  0.4815   0.4882
Day.Infected                 1   625.5  625.50 21.2401 5.68e-06 ***
Sex                          1     2.0    1.98  0.0673   0.7954
Infection.level:Day.Infected   1     1.6    1.59  0.0542   0.8161
Infection.level:Sex          1     8.7    8.65  0.2938   0.5881
Residuals                  351 10336.7   29.45
---
Signif. codes:  0 ‘***’ 0.001 ‘**’ 0.01 ‘*’ 0.05 ‘.’ 0.1 ‘ ’ 1
243 observations deleted due to missingness
> boxplot(Died~Day.Infected)
> ded4<-update(ded3,~.-Infection:level:Day.Infected)
> summary(ded4)
                            Df  Sum Sq Mean Sq F value   Pr(>F)
Infection.level              1    14.2   14.18  0.4815   0.4882
Day.Infected                 1   625.5  625.50 21.2401 5.68e-06 ***
Sex                          1     2.0    1.98  0.0673   0.7954
Infection.level:Day.Infected   1     1.6    1.59  0.0542   0.8161
Infection.level:Sex          1     8.7    8.65  0.2938   0.5881
Residuals                  351 10336.7   29.45
---
Signif. codes:  0 ‘***’ 0.001 ‘**’ 0.01 ‘*’ 0.05 ‘.’ 0.1 ‘ ’ 1
243 observations deleted due to missingness
> ded4<-update(ded3,~.-Infection.level:Day.Infected)
> summary(ded4)
                     Df  Sum Sq Mean Sq F value    Pr(>F)
Infection.level       1    14.2   14.18  0.4828    0.4876
Day.Infected          1   625.5  625.50 21.2975 5.517e-06 ***
Sex                   1     2.0    1.98  0.0675    0.7952
Infection.level:Sex   1     8.7    8.72  0.2969    0.5862
Residuals           352 10338.2   29.37
```

```
---
Signif. codes:  0 ‘***’ 0.001 ‘**’ 0.01 ‘*’ 0.05 ‘.’ 0.1 ‘ ’ 1
243 observations deleted due to missingness
> ded5<-update(ded4,~.-Infection.level:Sex)
> summary(ded5)
                 Df  Sum Sq Mean Sq F value    Pr(>F)
Infection.level   1    14.2   14.18  0.4838    0.4872
Day.Infected      1   625.5  625.50 21.3400 5.398e-06 ***
Sex               1     2.0    1.98  0.0676    0.7950
Residuals       353 10346.9   29.31
---
Signif. codes:  0 ‘***’ 0.001 ‘**’ 0.01 ‘*’ 0.05 ‘.’ 0.1 ‘ ’ 1
243 observations deleted due to missingness
> ded6<-update(ded5,~.-Sex)
> summary(ded6)
                 Df  Sum Sq Mean Sq F value    Pr(>F)
Infection.level   1    17.6   17.64  0.4074    0.5237
Day.Infected      1  1176.6 1176.58 27.1704 3.002e-07 ***
Residuals       397 17191.5   43.30
---
Signif. codes:  0 ‘***’ 0.001 ‘**’ 0.01 ‘*’ 0.05 ‘.’ 0.1 ‘ ’ 1
200 observations deleted due to missingness
> ded7<-aov(Died~DayInfected)
Error in eval(expr, envir, enclos) : object 'DayInfected' not found
> ded7<-aov(Died~Day.Infected)
> summary(ded7)
              Df  Sum Sq Mean Sq F value    Pr(>F)
Day.Infected   1  1176.6 1176.58  27.211 2.941e-07 ***
Residuals    398 17209.2   43.24
---
Signif. codes:  0 ‘***’ 0.001 ‘**’ 0.01 ‘*’ 0.05 ‘.’ 0.1 ‘ ’ 1
200 observations deleted due to missingness
> ded8<-lm(Died~Day.Infected)
> summary(ded8)

Call:
lm(formula = Died ~ Day.Infected)

Residuals:
    Min      1Q  Median      3Q     Max
-17.126  -4.541   1.408   4.476  16.476

Coefficients:
             Estimate Std. Error t value Pr(>|t|)
(Intercept)   16.7570     0.6738  24.869  < 2e-16 ***
Day.Infected   0.7670     0.1470   5.216 2.94e-07 ***
---
Signif. codes:  0 ‘***’ 0.001 ‘**’ 0.01 ‘*’ 0.05 ‘.’ 0.1 ‘ ’ 1

Residual standard error: 6.576 on 398 degrees of freedom
  (200 observations deleted due to missingness)
Multiple R-squared: 0.06399,    Adjusted R-squared: 0.06164
F-statistic: 27.21 on 1 and 398 DF,  p-value: 2.941e-07
```

```
> names(nan)
[1] "Plate"           "Infection.level" "Day.Infected"    "Pupated"
[5] "Died"            "Sex"             "Wing.length"     "Spore.count"
[9] "Duration"
> Ad<-(Died-Pupation)
Error: object 'Pupation' not found
> Ad<-(Died-Pupated)
> ad1<-aov(Ad~Infection.level*Day.Infected*Sex)
> summary(ad1)
                                 Df  Sum Sq Mean Sq F value    Pr(>F)
Infection.level                   1     9.6    9.63  0.3254    0.5687
Day.Infected                      1   559.0  558.98 18.8959 1.815e-05 ***
Sex                               1     0.5    0.54  0.0183    0.8924
Infection.level:Day.Infected      1     3.3    3.35  0.1132    0.7368
Infection.level:Sex               1     8.4    8.35  0.2824    0.5955
Day.Infected:Sex                  1     0.5    0.47  0.0158    0.9001
Infection.level:Day.Infected:Sex  1    51.9   51.89  1.7542    0.1862
Residuals                       348 10294.6   29.58
---
Signif. codes:  0 ‘***’ 0.001 ‘**’ 0.01 ‘*’ 0.05 ‘.’ 0.1 ‘ ’ 1
244 observations deleted due to missingness
> ad2<-aov(Ad~Infection.level+Day.Infected+Sex)
> anova(ad1,ad2)
Analysis of Variance Table

Model 1: Ad ~ Infection.level * Day.Infected * Sex
Model 2: Ad ~ Infection.level + Day.Infected + Sex
  Res.Df   RSS Df Sum of Sq      F Pr(>F)
1    348 10295
2    352 10359 -4   -64.061 0.5414 0.7054
> summary(ad2)
                 Df  Sum Sq Mean Sq F value    Pr(>F)
Infection.level   1     9.6    9.63  0.3271    0.5677
Day.Infected      1   559.0  558.98 18.9949 1.722e-05 ***
Sex               1     0.5    0.54  0.0184    0.8921
Residuals       352 10358.6   29.43
---
Signif. codes:  0 ‘***’ 0.001 ‘**’ 0.01 ‘*’ 0.05 ‘.’ 0.1 ‘ ’ 1
244 observations deleted due to missingness
> ad3<-aov(Ad~Infection.level+Day.Infected)
> ad4<-aov(Ad~Day.Infected)
> anova(ad2,ad3,ad4)
Error in anova.lmlist(object, ...) :
  models were not all fitted to the same size of dataset
> anova(ad2,ad3)
Error in anova.lmlist(object, ...) :
  models were not all fitted to the same size of dataset
> summary(ad4)
              Df  Sum Sq Mean Sq F value    Pr(>F)
Day.Infected   1   439.2  439.17  12.707 0.0004113 ***
Residuals    376 12995.3   34.56
---
Signif. codes:  0 ‘***’ 0.001 ‘**’ 0.01 ‘*’ 0.05 ‘.’ 0.1 ‘ ’ 1
222 observations deleted due to missingness
```

```
> ad5<-lm(Ad~Day.Infected)
> summary(ad5)

Call:
lm(formula = Ad ~ Day.Infected)

Residuals:
    Min      1Q  Median      3Q     Max 
-13.839  -3.926   1.103   4.103  15.074 

Coefficients:
             Estimate Std. Error t value Pr(>|t|)    
(Intercept)   11.4401     0.6339  18.049  < 2e-16 ***
Day.Infected   0.4855     0.1362   3.565 0.000411 ***
---
Signif. codes:  0 ‘***’ 0.001 ‘**’ 0.01 ‘*’ 0.05 ‘.’ 0.1 ‘ ’ 1

Residual standard error: 5.879 on 376 degrees of freedom
  (222 observations deleted due to missingness)
Multiple R-squared: 0.03269,    Adjusted R-squared: 0.03012 
F-statistic: 12.71 on 1 and 376 DF,  p-value: 0.0004113

>
```
