## Supplementary material for "*Anopheles coluzzii* infection by the microsporidian, *Vavraia culicis*: the effect of host age": Statistical analysis in R (2)

[Day.Infected!=NA]

a maximal quasibinomial glm was fit to the data. Spore count was the response variable, Day infected, sex, infection level were the explanatory variables.
Model simplification was carried out by stepwise elimination of non-significant factors and interactions.

There were no significant interactions between the explanatory variables.
The original level of infection did not affect the spore count at the end of the mosquito's life (t=0.501, d.f.=1, p=0.62).
Male mosquitoes produced fewer spores than did females (t=3.31, d.f.=1, p<0.01).
The day infected also affected spore production; later infections produced fewer spores (t=9.45, d.f.=1, p<0.001).
Even when the duration of infection was the same, later infections still produced fewer spores compared to earlier infections.

Infection does not affect the age at death of the mosquitoes (t = 0.77, df = 598, p-value = 0.44)

```
R version 2.7.2 (2008-08-25)
Copyright (C) 2008 The R Foundation for Statistical Computing
ISBN 3-900051-07-0

R is free software and comes with ABSOLUTELY NO WARRANTY.
You are welcome to redistribute it under certain conditions.
Type 'license()' or 'licence()' for distribution details.

  Natural language support but running in an English locale

R is a collaborative project with many contributors.
Type 'contributors()' for more information and
'citation()' on how to cite R or R packages in publications.

Type 'demo()' for some demos, 'help()' for on-line help, or
'help.start()' for an HTML browser interface to help.
Type 'q()' to quit R.

[Previously saved workspace restored]

> nan<-read.table("D://Nayna.txt",header=T)
> names(nan)
[1] "Plate"           "Infection.level" "Day.Infected"    "Pupated"
[5] "Died"            "Sex"             "Wing.length"     "Spore.count"
[9] "Duration"
> attach(nan)
> plot(Day.Infected,Duration)
> summary(Day.infected)
Error in summary(Day.infected) : object "Day.infected" not found
> summary(Day.Infected)
   Min. 1st Qu.  Median    Mean 3rd Qu.    Max.    NA's
    1.0     2.5     4.0     4.0     5.5     7.0   200.0
> hist(Spore.Count)
Error in hist(Spore.Count) : object "Spore.Count" not found
> hist(Spore.count)
```

```
> hist(log(Spore.count))
> q11<-glm(Spore.count~DayInfected*Sex*Infection.level*Duration+Plate,poisson)
Error in eval(expr, envir, enclos) : object "DayInfected" not found
> q11<-glm(Spore.count~Day.Infected*Sex*Infection.level*Duration+Plate,poisson)
Error: NA/NaN/Inf in foreign function call (arg 1)
In addition: Warning message:
step size truncated due to divergence
> summary(q11)
Error in summary(q11) : object "q11" not found
> q11<-glm(Spore.count~Day.Infected*Sex*Infection.level+Duration+Plate,poisson)
Error: NA/NaN/Inf in foreign function call (arg 1)
In addition: Warning message:
step size truncated due to divergence
> q11<-glm(Spore.count~Day.Infected*Sex+Infection.level+Duration+Plate,poisson)
Error: NA/NaN/Inf in foreign function call (arg 1)
In addition: Warning message:
step size truncated due to divergence
> q11<-glm(Spore.count~Day.Infected+Sex+Infection.level+Duration+Plate,poisson)
Error: NA/NaN/Inf in foreign function call (arg 1)
In addition: Warning message:
step size truncated due to divergence
> q11<-
glm(Spore.count[Day.Infected1=!NA]~Day.Infected[Day.Infected!=NA]+Sex[Day.Infected!=NA]+Infection.level[Day.
Infected!=NA]+Duration[Day.Infected!=NA]+Plate[Day.Infected!=NA],poisson)
Error in `contrasts<-`(`*tmp*`, value = "contr.treatment") :
  contrasts can be applied only to factors with 2 or more levels
> summary(q11)
Error in summary(q11) : object "q11" not found
> q11<-glm(Spore.count[Day.Infected1=!NA]~Day.Infected[Day.Infected!=NA],poisson)
Error in pmax(exp(eta), .Machine$double.eps) :
  cannot mix 0-length vectors with others
> q11<-glm(Spore.count[Day.Infected!=NA]~Day.Infected[Day.Infected!=NA],poisson)
Error in pmax(exp(eta), .Machine$double.eps) :
  cannot mix 0-length vectors with others
> q11<-glm(Spore.count~Day.Infected,poisson)
> summary(q11)

Call:
glm(formula = Spore.count ~ Day.Infected, family = poisson)

Deviance Residuals:
     Min        1Q    Median        3Q       Max
-1024.31   -422.03   -261.40     50.33   2964.10

Coefficients:
              Estimate Std. Error z value Pr(>|z|)
(Intercept)   1.363e+01  1.764e-04   77249   <2e-16 ***
Day.Infected -4.552e-01  6.538e-05   -6963   <2e-16 ***
---
Signif. codes:  0 ‘***’ 0.001 ‘**’ 0.01 ‘*’ 0.05 ‘.’ 0.1 ‘ ’ 1

(Dispersion parameter for poisson family taken to be 1)

    Null deviance: 233045876  on 399  degrees of freedom
```

```
Residual deviance: 167534374  on 398  degrees of freedom
  (200 observations deleted due to missingness)
AIC: 167537203

Number of Fisher Scoring iterations: 6

> boxplot(Spore.count~Day.Infected)
> q12<-glm(Spore.count~Day.Infected,quasipoisson)
> summary(q12)

Call:
glm(formula = Spore.count ~ Day.Infected, family = quasipoisson)

Deviance Residuals: 
     Min        1Q    Median        3Q       Max  
-1024.31   -422.03   -261.40     50.33   2964.10  

Coefficients:
             Estimate Std. Error t value Pr(>|t|)    
(Intercept)  13.62565    0.14072  96.825   <2e-16 ***
Day.Infected -0.45524    0.05216  -8.728   <2e-16 ***
---
Signif. codes:  0 ‘***’ 0.001 ‘**’ 0.01 ‘*’ 0.05 ‘.’ 0.1 ‘ ’ 1

(Dispersion parameter for quasipoisson family taken to be 636523.6)

    Null deviance: 233045876  on 399  degrees of freedom
Residual deviance: 167534374  on 398  degrees of freedom
  (200 observations deleted due to missingness)
AIC: NA

Number of Fisher Scoring iterations: 6


> q14<-glm(Spore.count~Day.Infected*Sex*Infection.level,quasipoisson)
> summary(q14)

Call:
glm(formula = Spore.count ~ Day.Infected * Sex * Infection.level, 
    family = quasipoisson)

Deviance Residuals: 
     Min        1Q    Median        3Q       Max  
-1298.46   -417.49   -219.66     71.17   2593.66  

Coefficients:
                                 Estimate Std. Error t value Pr(>|t|)    
(Intercept)                      14.25261    0.22463  63.451  < 2e-16 ***
Day.Infected                     -0.60789    0.09645  -6.303 8.83e-10 ***
SexM                             -1.26203    0.43936  -2.872  0.00432 ** 
Infection.levellow               -0.35708    0.31567  -1.131  0.25877    
Day.Infected:SexM                 0.29027    0.14912   1.947  0.05239 .  
Day.Infected:Infection.levellow   0.20346    0.12349   1.648  0.10034    
SexM:Infection.levellow           0.90639    0.56955   1.591  0.11242    
```

```
Day.Infected:SexM:Infection.levellow -0.38494    0.20410  -1.886  0.06012 .
---
Signif. codes:  0 ‘***’ 0.001 ‘**’ 0.01 ‘*’ 0.05 ‘.’ 0.1 ‘ ’ 1

(Dispersion parameter for quasipoisson family taken to be 526653.3)

    Null deviance: 213603944  on 356  degrees of freedom
Residual deviance: 136475259  on 349  degrees of freedom
  (243 observations deleted due to missingness)
AIC: NA

Number of Fisher Scoring iterations: 7

> q15<-update(q14,~.-Day.Infected:Sex:Infection.level)
> Anova(q14,q15,Test="Chi")
Error: could not find function "Anova"
> anova(q14,q15,Test="Chi")
Analysis of Deviance Table

Model 1: Spore.count ~ Day.Infected * Sex * Infection.level
Model 2: Spore.count ~ Day.Infected + Sex + Infection.level + Day.Infected:Sex +
    Day.Infected:Infection.level + Sex:Infection.level
  Resid. Df Resid. Dev  Df  Deviance
1       349  136475259
2       350  138374856  -1  -1899597
Warning message:
In anova.glm(q14, q15, Test = "Chi") :
  the following arguments to 'anova.glm' are invalid and dropped: structure(list(Test = "Chi"), .Names = "Test")
> anova(q14,q15,Test="F")
Analysis of Deviance Table

Model 1: Spore.count ~ Day.Infected * Sex * Infection.level
Model 2: Spore.count ~ Day.Infected + Sex + Infection.level + Day.Infected:Sex +
    Day.Infected:Infection.level + Sex:Infection.level
  Resid. Df Resid. Dev  Df  Deviance
1       349  136475259
2       350  138374856  -1  -1899597
Warning message:
In anova.glm(q14, q15, Test = "F") :
  the following arguments to 'anova.glm' are invalid and dropped: structure(list(Test = "F"), .Names = "Test")
> summary(q15)

Call:
glm(formula = Spore.count ~ Day.Infected + Sex + Infection.level +
    Day.Infected:Sex + Day.Infected:Infection.level + Sex:Infection.level,
    family = quasipoisson)

Deviance Residuals:
    Min       1Q   Median       3Q      Max
-1250.9   -434.8   -272.1     78.9   2638.4

Coefficients:
                             Estimate Std. Error t value Pr(>|t|)
(Intercept)                  14.09841    0.21691  64.997  < 2e-16 ***
```

```
Day.Infected                   -0.52840    0.08176  -6.463 3.45e-10 ***
SexM                           -0.76900    0.34819  -2.209   0.0278 *
Infection.levellow             -0.07163    0.28563  -0.251   0.8021
Day.Infected:SexM               0.07411    0.10306   0.719   0.4726
Day.Infected:Infection.levellow  0.06455    0.10000   0.645   0.5191
SexM:Infection.levellow         0.05587    0.35145   0.159   0.8738
---
Signif. codes:  0 '***' 0.001 '**' 0.01 '*' 0.05 '.' 0.1 ' ' 1

(Dispersion parameter for quasipoisson family taken to be 558933.6)

    Null deviance: 213603944  on 356  degrees of freedom
Residual deviance: 138374856  on 350  degrees of freedom
  (243 observations deleted due to missingness)
AIC: NA

Number of Fisher Scoring iterations: 7

> q16<-update(q15,~.-Sex:Infection.level)
> anova(q14,q15,Test="F")
Analysis of Deviance Table

Model 1: Spore.count ~ Day.Infected * Sex * Infection.level
Model 2: Spore.count ~ Day.Infected + Sex + Infection.level + Day.Infected:Sex +
    Day.Infected:Infection.level + Sex:Infection.level
  Resid. Df Resid. Dev  Df  Deviance
1       349  136475259
2       350  138374856  -1  -1899597
Warning message:
In anova.glm(q14, q15, Test = "F") :
  the following arguments to 'anova.glm' are invalid and dropped: structure(list(Test = "F"), .Names = "Test")
> anova(q16,q15,Test="Chi")
Analysis of Deviance Table

Model 1: Spore.count ~ Day.Infected + Sex + Infection.level + Day.Infected:Sex +
    Day.Infected:Infection.level
Model 2: Spore.count ~ Day.Infected + Sex + Infection.level + Day.Infected:Sex +
    Day.Infected:Infection.level + Sex:Infection.level
  Resid. Df Resid. Dev  Df  Deviance
1       351  138389002
2       350  138374856   1     14145
Warning message:
In anova.glm(q16, q15, Test = "Chi") :
  the following arguments to 'anova.glm' are invalid and dropped: structure(list(Test = "Chi"), .Names = "Test")
> summary(q16)

Call:
glm(formula = Spore.count ~ Day.Infected + Sex + Infection.level +
    Day.Infected:Sex + Day.Infected:Infection.level, family = quasipoisson)

Deviance Residuals:
     Min        1Q    Median        3Q       Max
-1251.64   -432.44   -268.42     75.59   2654.26
```

```
Coefficients:
                               Estimate Std. Error t value Pr(>|t|)
(Intercept)                    14.09073    0.21164  66.578  < 2e-16 ***
Day.Infected                   -0.52900    0.08174  -6.472 3.27e-10 ***
SexM                           -0.73735    0.28389  -2.597   0.0098 **
Infection.levellow             -0.05581    0.26684  -0.209   0.8344
Day.Infected:SexM               0.07410    0.10272   0.721   0.4711
Day.Infected:Infection.levellow  0.06535    0.09953   0.657   0.5119
---
Signif. codes:  0 '***' 0.001 '**' 0.01 '*' 0.05 '.' 0.1 ' ' 1

(Dispersion parameter for quasipoisson family taken to be 556098.2)

    Null deviance: 213603944  on 356  degrees of freedom
Residual deviance: 138389002  on 351  degrees of freedom
  (243 observations deleted due to missingness)
AIC: NA

Number of Fisher Scoring iterations: 6

> q17<-update(q16,~.-Day.Infected:Infection.level)
> summary(q17)

Call:
glm(formula = Spore.count ~ Day.Infected + Sex + Infection.level + 
    Day.Infected:Sex, family = quasipoisson)

Deviance Residuals: 
     Min        1Q    Median        3Q       Max  
-1274.31   -454.11   -261.40     80.74   2652.84  

Coefficients:
                  Estimate Std. Error t value Pr(>|t|)
(Intercept)       14.01833    0.18343  76.422  < 2e-16 ***
Day.Infected      -0.49496    0.06206  -7.975 2.16e-14 ***
SexM              -0.74979    0.28478  -2.633  0.00884 **
Infection.levellow  0.08381    0.16299   0.514  0.60743
Day.Infected:SexM   0.07876    0.10308   0.764  0.44533
---
Signif. codes:  0 '***' 0.001 '**' 0.01 '*' 0.05 '.' 0.1 ' ' 1

(Dispersion parameter for quasipoisson family taken to be 561361.3)

    Null deviance: 213603944  on 356  degrees of freedom
Residual deviance: 138629871  on 352  degrees of freedom
  (243 observations deleted due to missingness)
AIC: NA

Number of Fisher Scoring iterations: 6

> anova(q16,q17,Test="Chi")
Analysis of Deviance Table

Model 1: Spore.count ~ Day.Infected + Sex + Infection.level + Day.Infected:Sex + 
```

```
    Day.Infected:Infection.level
Model 2: Spore.count ~ Day.Infected + Sex + Infection.level + Day.Infected:Sex
  Resid. Df Resid. Dev  Df  Deviance
1       351  138389002
2       352  138629871  -1   -240870
Warning message:
In anova.glm(q16, q17, Test = "Chi") :
  the following arguments to 'anova.glm' are invalid and dropped: structure(list(Test = "Chi"), .Names = "Test")
> q18<-update(q17,~.-Day.Infected:Sex)
> summary(q18)

Call:
glm(formula = Spore.count ~ Day.Infected + Sex + Infection.level,
    family = quasipoisson)

Deviance Residuals:
     Min        1Q    Median        3Q       Max
-1254.59   -472.41   -251.31     80.22   2632.31

Coefficients:
                   Estimate Std. Error t value Pr(>|t|)
(Intercept)        13.96273    0.16833  82.948  < 2e-16 ***
Day.Infected       -0.46803    0.04938  -9.478  < 2e-16 ***
SexM               -0.58037    0.17322  -3.350 0.000894 ***
Infection.levellow  0.08129    0.16221   0.501 0.616578
---
Signif. codes:  0 ‘***’ 0.001 ‘**’ 0.01 ‘*’ 0.05 ‘.’ 0.1 ‘ ’ 1

(Dispersion parameter for quasipoisson family taken to be 555963.8)

    Null deviance: 213603944  on 356  degrees of freedom
Residual deviance: 138951779  on 353  degrees of freedom
  (243 observations deleted due to missingness)
AIC: NA

Number of Fisher Scoring iterations: 6

> q19<-update(q18,~.-Infection.level)
> summary(q19)

Call:
glm(formula = Spore.count ~ Day.Infected + Sex, family = quasipoisson)

Deviance Residuals:
     Min        1Q    Median        3Q       Max
-1230.04   -481.53   -258.34     91.43   2669.66

Coefficients:
             Estimate Std. Error t value Pr(>|t|)
(Intercept)   14.0054     0.1449  96.643  < 2e-16 ***
Day.Infected  -0.4689     0.0496  -9.454  < 2e-16 ***
SexM          -0.5743     0.1736  -3.309  0.00103 **
---
Signif. codes:  0 ‘***’ 0.001 ‘**’ 0.01 ‘*’ 0.05 ‘.’ 0.1 ‘ ’ 1
```

```
(Dispersion parameter for quasipoisson family taken to be 560912.5)

    Null deviance: 213603944  on 356  degrees of freedom
Residual deviance: 139091634  on 354  degrees of freedom
  (243 observations deleted due to missingness)
AIC: NA

Number of Fisher Scoring iterations: 6

> anova(q19,q18,Test="Chi")
Analysis of Deviance Table

Model 1: Spore.count ~ Day.Infected + Sex
Model 2: Spore.count ~ Day.Infected + Sex + Infection.level
  Resid. Df Resid. Dev  Df  Deviance
1       354  139091634
2       353  138951779   1    139855
Warning message:
In anova.glm(q19, q18, Test = "Chi") :
  the following arguments to 'anova.glm' are invalid and dropped: structure(list(Test = "Chi"), .Names = "Test")
> names(nan)
[1] "Plate"           "Infection.level" "Day.Infected"    "Pupated"         "Died"
[6] "Sex"             "Wing.length"     "Spore.count"     "Duration"
> hist(died)
Error in hist(died) : object "died" not found
> hist(Died)
> plot(Died,Day.Infected)
> plot(Died~Day.Infected)

> length(Day.Infected)
[1] 600
> 600/12
[1] 50
> I<-rep(c(yes,yes,yes,yes,yes,yes,yes,yes,no,no,no,no),50)
Error: object "yes" not found
> I<-rep(c("yes","yes","yes","yes","yes","yes","yes","yes","no","no","no","no"),50)

> tapply(Died,I,mean)
    no    yes
20.295 19.825
> var.test(Died~I)

        F test to compare two variances

data:  Died by I
F = 1.2026, num df = 199, denom df = 399, p-value = 0.1260
alternative hypothesis: true ratio of variances is not equal to 1
95 percent confidence interval:
 0.9496114 1.5380428
sample estimates:
ratio of variances
          1.202595
```

```
> t.test(Died~I)

        Welch Two Sample t-test

data:  Died by I
t = 0.7504, df = 367.216, p-value = 0.4535
alternative hypothesis: true difference in means is not equal to 0
95 percent confidence interval:
 -0.7616199  1.7016199
sample estimates:
 mean in group no mean in group yes
           20.295            19.825

> t.test(Died~I,var.equal=T)

        Two Sample t-test

data:  Died by I
t = 0.7738, df = 598, p-value = 0.4393
alternative hypothesis: true difference in means is not equal to 0
95 percent confidence interval:
 -0.7228346  1.6628346
sample estimates:
 mean in group no mean in group yes
           20.295            19.825

> death1<-glm(Died~I*Sex*Day.Infected)
Error in `contrasts<-`(`*tmp*`, value = "contr.treatment") :
  contrasts can be applied only to factors with 2 or more levels
In addition: Warning message:
In model.matrix.default(mt, mf, contrasts) :
  variable 'I' converted to a factor
> summary(death1)
Error in summary(death1) : object "death1" not found
> death1<-glm(Died~I*Sex*Day.Infected)
Error in `contrasts<-`(`*tmp*`, value = "contr.treatment") :
  contrasts can be applied only to factors with 2 or more levels
In addition: Warning message:
In model.matrix.default(mt, mf, contrasts) :
  variable 'I' converted to a factor
> is.factor(I)
[1] FALSE
> inf<-as.factor(I)
> death1<-glm(Died~inf*Sex*Day.Infected)
Error in `contrasts<-`(`*tmp*`, value = "contr.treatment") :
  contrasts can be applied only to factors with 2 or more levels
> death1<-glm(Died~inf*Sex)
> summary(death1)

Call:
glm(formula = Died ~ inf * Sex)

Deviance Residuals:
     Min        1Q    Median        3Q       Max
```

```
-12.5833   -3.2308    0.5631    3.7692   14.9198

Coefficients:
            Estimate Std. Error t value Pr(>|t|)
(Intercept)  21.4369     0.5744  37.320   <2e-16 ***
infyes       -0.2061     0.7101  -0.290    0.772
SexM          1.1464     0.8955   1.280    0.201
infyes:SexM  -1.2970     1.0890  -1.191    0.234
---
Signif. codes:  0 ‘***’ 0.001 ‘**’ 0.01 ‘*’ 0.05 ‘.’ 0.1 ‘ ’ 1

(Dispersion parameter for gaussian family taken to be 33.98373)

    Null deviance: 18066  on 531  degrees of freedom
Residual deviance: 17943  on 528  degrees of freedom
  (68 observations deleted due to missingness)
AIC: 3391.5

Number of Fisher Scoring iterations: 2

> death2<-lm(Died~inf*Sex*Day.Infected)
Error in `contrasts<-`(`*tmp*`, value = "contr.treatment") :
  contrasts can be applied only to factors with 2 or more levels
> death2<-lm(Died~inf*Sex)
> summary(death2)

Call:
lm(formula = Died ~ inf * Sex)

Residuals:
     Min       1Q   Median       3Q      Max
-12.5833  -3.2308   0.5631   3.7692  14.9198

Coefficients:
            Estimate Std. Error t value Pr(>|t|)
(Intercept)  21.4369     0.5744  37.320   <2e-16 ***
infyes       -0.2061     0.7101  -0.290    0.772
SexM          1.1464     0.8955   1.280    0.201
infyes:SexM  -1.2970     1.0890  -1.191    0.234
---
Signif. codes:  0 ‘***’ 0.001 ‘**’ 0.01 ‘*’ 0.05 ‘.’ 0.1 ‘ ’ 1

Residual standard error: 5.83 on 528 degrees of freedom
  (68 observations deleted due to missingness)
Multiple R-squared: 0.006812,   Adjusted R-squared: 0.001169
F-statistic: 1.207 on 3 and 528 DF,  p-value: 0.3064

> death3<-lm(Died~inf+Sex)
> summary(death3)

Call:
lm(formula = Died ~ inf + Sex)

Residuals:
```

```
     Min       1Q   Median       3Q      Max 
-12.3097  -3.0402   0.6903   3.9598  14.6903 

Coefficients:
            Estimate Std. Error t value Pr(>|t|)    
(Intercept)  21.7977     0.4882  44.649   <2e-16 ***
infyes       -0.7575     0.5386  -1.406    0.160    
SexM          0.2695     0.5098   0.529    0.597    
---
Signif. codes:  0 ‘***’ 0.001 ‘**’ 0.01 ‘*’ 0.05 ‘.’ 0.1 ‘ ’ 1 

Residual standard error: 5.832 on 529 degrees of freedom
  (68 observations deleted due to missingness)
Multiple R-squared: 0.004144,   Adjusted R-squared: 0.0003794 
F-statistic: 1.101 on 2 and 529 DF,  p-value: 0.3334 

> death4<-lm(Died~inf)
> summary(death4)

Call:
lm(formula = Died ~ inf)

Residuals:
    Min      1Q  Median      3Q     Max 
-15.825  -5.295   1.175   5.175  16.175 

Coefficients:
            Estimate Std. Error t value Pr(>|t|)    
(Intercept)  20.2950     0.4959  40.924   <2e-16 ***
infyes       -0.4700     0.6074  -0.774    0.439    
---
Signif. codes:  0 ‘***’ 0.001 ‘**’ 0.01 ‘*’ 0.05 ‘.’ 0.1 ‘ ’ 1 

Residual standard error: 7.013 on 598 degrees of freedom
Multiple R-squared: 0.001,      Adjusted R-squared: -0.0006702 
F-statistic: 0.5988 on 1 and 598 DF,  p-value: 0.4393 

> barplot(tapply(Spore.count,list(Sex,DayInfected),mean),beside=T)
Error in tapply(Spore.count, list(Sex, DayInfected), mean) : 
  object "DayInfected" not found
> barplot(tapply(Spore.count,list(Sex,Day.Infected),mean),beside=T)
> barplot(tapply(log(Spore.count),list(Sex,Day.Infected),mean),beside=T)
Error in plot.window(xlim, ylim, log = log, ...) : 
  need finite 'ylim' values
> barplot(tapply(Spore.count,list(Sex,Day.Infected),mean),beside=T)
> barplot(tapply(Spore.count,list(Sex,Day.Infected),mean),beside=T,ylab="Number of Spore produced",xlab="Day of inefction")
> 
```
