## Supplementary material for "*Anopheles coluzzii* infection by the microsporidian, *Vavraia culicis*: the effect of host age": Raw data for the experiment

| Plate | Infection.le | Day.Infecte | Pupated | Died | Sex | Wing.lengt | Spore.cour | Duration |
|---|---|---|---|---|---|---|---|---|
| 10A1 | low | 1 | 7 | 20 | M | NA | 3050000 | 19 |
| 10A2 | low | 3 | 7 | 25 | M | NA | 760000 | 22 |
| 10A3 | low | 5 | 7 | 26 | M | NA | 20000 | 21 |
| 10A4 | low | 7 | 7 | 21 | F | 3.224 | 0 | NA |
| 10B1 | high | 1 | 7 | 24 | F | 3.189 | 500000 | 23 |
| 10B2 | high | 3 | 7 | 25 | F | 3.404 | 450000 | 22 |
| 10B3 | high | 5 | 7 | 21 | M | NA | 340000 | 16 |
| 10B4 | high | 7 | 7 | 26 | F | 3.352 | 0 | NA |
| 10C1 | control | NA | 8 | 20 | F | 3.652 | 0 | NA |
| 10C2 | control | NA | 7 | 24 | F | 3.444 | 0 | NA |
| 10C3 | control | NA | 7 | 23 | M | NA | 0 | NA |
| 10C4 | control | NA | 7 | 26 | F | 3.517 | 0 | NA |
| 11A1 | low | 1 | 7 | 9 | M | 3.368 | 0 | 8 |
| 11A2 | low | 3 | 7 | 23 | M | NA | 300000 | 20 |
| 11A3 | low | 5 | 7 | 28 | F | 3.3 | 60000 | 23 |
| 11A4 | low | 7 | 7 | 22 | F | 3.27 | 0 | NA |
| 11B1 | high | 1 | 7 | 21 | M | NA | 50000 | 20 |
| 11B2 | high | 3 | 8 | 22 | F | NA | 350000 | 19 |
| 11B3 | high | 5 | 7 | 25 | F | 3.516 | 0 | 20 |
| 11B4 | high | 7 | 8 | 13 | F | 3.152 | 0 | NA |
| 11C1 | control | NA | 8 | 23 | M | NA | 0 | NA |
| 11C2 | control | NA | 9 | 26 | F | 3.184 | 0 | NA |
| 11C3 | control | NA | 8 | 21 | M | NA | 0 | NA |
| 11C4 | control | NA | 8 | 22 | M | NA | 0 | NA |
| 12A1 | low | 1 | 7 | 10 | F | 3.745 | 0 | 9 |
| 12A2 | low | 3 | 7 | 21 | F | 3.308 | 300000 | 18 |
| 12A3 | low | 5 | 7 | 12 | M | NA | 0 | 7 |
| 12A4 | low | 7 | 7 | 27 | F | 3.278 | 13000 | 20 |
| 12B1 | high | 1 | 7 | 21 | M | NA | 0 | 20 |
| 12B2 | high | 3 | 7 | 22 | M | NA | 27000 | 19 |
| 12B3 | high | 5 | 7 | 22 | F | 3.53 | 0 | NA |
| 12B4 | high | 7 | 8 | 27 | F | 3.548 | 0 | NA |
| 12C1 | control | NA | 7 | 22 | F | 3.272 | 0 | NA |
| 12C2 | control | NA | 7 | 25 | M | NA | 0 | NA |
| 12C3 | control | NA | 7 | 25 | M | NA | 0 | NA |
| 12C4 | control | NA | NA | 6 | NA | NA | 0 | NA |
| 13A1 | low | 1 | 7 | 22 | M | NA | 920000 | 21 |
| 13A2 | low | 3 | 7 | 13 | NA | NA | 0 | 10 |
| 13A3 | low | 5 | 7 | 18 | M | NA | 20000 | 16 |
| 13A4 | low | 7 | 7 | 23 | M | NA | 0 | na |
| 13B1 | high | 1 | 7 | 18 | F | 3.23 | 1190000 | 17 |
| 13B2 | high | 3 | 7 | 15 | M | NA | 20000 | 12 |
| 13B3 | high | 5 | 9 | 21 | M | NA | 50000 | 16 |
| 13B4 | high | 7 | 7 | 16 | F | 3.308 | 0 | NA |
| 13C1 | control | NA | NA | 7 | NA | NA | 0 | NA |
| 13C2 | control | NA | 7 | 9 | NA | NA | 0 | NA |
| 13C3 | control | NA | 7 | 34 | M | NA | 0 | NA |
| 13C4 | control | NA | 7 | 23 | F | 3.485 | 0 | NA |
| 14A1 | low | 1 | 7 | 26 | M | NA | 0 | 19 |
| 14A2 | low | 3 | 7 | 21 | M | NA | 40000 | 18 |

| | | | | | | | | |
|---|---|---|---|---|---|---|---|---|
| 14A3 | low | 5 | 7 | 21 | M | NA | 170000 | 16 |
| 14A4 | low | 7 | 7 | 15 | M | NA | 0 | 8 |
| 14B1 | high | 1 | 7 | 26 | M | NA | 620000 | 25 |
| 14B2 | high | 3 | 7 | 12 | M | NA | 0 | 9 |
| 14B3 | high | 5 | 7 | 16 | F | 3.343 | 0 | 11 |
| 14B4 | high | 7 | 8 | 22 | F | 3.334 | 180000 | 15 |
| 14C1 | control | NA | 7 | 22 | F | 3.34 | 0 | NA |
| 14C2 | control | NA | 6 | 7 | NA | NA | 0 | NA |
| 14C3 | control | NA | 7 | 21 | M | NA | 0 | NA |
| 14C4 | control | NA | 7 | 22 | M | NA | 0 | NA |
| 15A1 | low | 1 | 7 | 25 | M | NA | 740000 | 24 |
| 15A2 | low | 3 | 9 | 13 | F | 3.263 | 10000 | 10 |
| 15A3 | low | 5 | 8 | 9 | NA | NA | 0 | 4 |
| 15A4 | low | 7 | 8 | 22 | M | NA | 0 | NA |
| 15B1 | high | 1 | 7 | 21 | F | 3.665 | 1360000 | 20 |
| 15B2 | high | 3 | 9 | 32 | F | NA | 240000 | 29 |
| 15B3 | high | 5 | 7 | 22 | M | NA | 0 | 17 |
| 15B4 | high | 7 | 7 | 21 | M | NA | 0 | NA |
| 15C1 | control | NA | 7 | 29 | M | NA | 0 | NA |
| 15C2 | control | NA | 8 | 19 | F | 3.671 | 0 | NA |
| 15C3 | control | NA | 7 | 25 | F | 2.985 | 0 | NA |
| 15C4 | control | NA | 7 | 22 | M | NA | 0 | NA |
| 16A1 | low | 1 | 7 | 26 | F | 3.374 | 660000 | 25 |
| 16A2 | low | 3 | 7 | 21 | F | 3.575 | 400000 | 18 |
| 16A3 | low | 5 | 9 | 15 | M | NA | 30000 | 10 |
| 16A4 | low | 7 | 9 | 26 | F | 3.229 | 0 | NA |
| 16B1 | high | 1 | 7 | 13 | F | 3.55 | 20000 | 12 |
| 16B2 | high | 3 | 7 | 13 | M | NA | 0 | 10 |
| 16B3 | high | 5 | 9 | 22 | F | 3.233 | 0 | 17 |
| 16B4 | high | 7 | 7 | 15 | M | NA | 0 | NA |
| 16C1 | control | NA | 9 | 15 | M | NA | 0 | NA |
| 16C2 | control | NA | 7 | 23 | F | 3.085 | 0 | NA |
| 16C3 | control | NA | 8 | 15 | F | 3.381 | 0 | NA |
| 16C4 | control | NA | 8 | 32 | M | NA | 0 | NA |
| 17A1 | low | 1 | 7 | 34 | F | 2.936 | 1100000 | 33 |
| 17A2 | low | 3 | 7 | 21 | F | 3.444 | 200000 | 18 |
| 17A3 | low | 5 | 7 | 21 | F | 3.353 | 200000 | 16 |
| 17A4 | low | 7 | 7 | 24 | M | NA | 0 | NA |
| 17B1 | high | 1 | 7 | 23 | M | NA | 0 | 22 |
| 17B2 | high | 3 | 8 | 14 | M | NA | 0 | 11 |
| 17B3 | high | 5 | 7 | 9 | M | NA | 0 | 4 |
| 17B4 | high | 7 | 7 | 26 | M | NA | 0 | NA |
| 17C1 | control | NA | 7 | 14 | F | 2.45 | 0 | NA |
| 17C2 | control | NA | 7 | 16 | F | 3.429 | 0 | NA |
| 17C3 | control | NA | NA | 9 | NA | NA | 0 | NA |
| 17C4 | control | NA | 7 | 21 | M | NA | 0 | NA |
| 18A1 | low | 1 | 7 | 22 | F | 3.642 | 440000 | 21 |
| 18A2 | low | 3 | 7 | 26 | M | NA | 350000 | 23 |
| 18A3 | low | 5 | 7 | 24 | F | 3.34 | 230000 | 19 |
| 18A4 | low | 7 | 8 | 29 | M | NA | 0 | NA |
| 18B1 | high | 1 | 7 | 22 | F | 3.34 | 2670000 | 21 |

| | | | | | | | | |
|---|---|---|---|---|---|---|---|---|
| 18B2 | high | 3 | 8 | 14 | F | NA | 0 | 11 |
| 18B3 | high | 5 | 7 | 21 | F | 2.55 | 0 | 16 |
| 18B4 | high | 7 | 7 | 36 | M | NA | 0 | NA |
| 18C1 | control | NA | 7 | 26 | M | NA | 0 | NA |
| 18C2 | control | NA | 7 | 19 | F | 3.28 | 0 | NA |
| 18C3 | control | NA | 7 | 10 | F | 3.013 | 0 | NA |
| 18C4 | control | NA | 8 | 10 | NA | NA | 0 | NA |
| 19A1 | low | 1 | 7 | 23 | M | NA | 0 | 22 |
| 19A2 | low | 3 | 7 | 24 | F | 3.523 | 770000 | 21 |
| 19A3 | low | 5 | 8 | 26 | M | NA | 30000 | 21 |
| 19A4 | low | 7 | 7 | 30 | M | NA | 0 | NA |
| 19B1 | high | 1 | NA | 4 | NA | NA | 0 | 3 |
| 19B2 | high | 3 | 7 | 22 | F | 3.464 | 280000 | 19 |
| 19B3 | high | 5 | 7 | 9 | NA | NA | 0 | 4 |
| 19B4 | high | 7 | 7 | 22 | M | NA | 0 | NA |
| 19C1 | control | NA | 7 | 12 | M | NA | 0 | NA |
| 19C2 | control | NA | 7 | 21 | F | 3.324 | 0 | NA |
| 19C3 | control | NA | 8 | 21 | M | NA | 0 | NA |
| 19C4 | control | NA | 7 | 25 | F | 3.327 | 0 | NA |
| 1A1 | low | 1 | 7 | 15 | M | NA | 0 | 14 |
| 1A2 | low | 3 | 7 | 14 | F | 3.261 | 0 | 11 |
| 1A3 | low | 5 | 7 | 21 | F | 3.605 | 70000 | 16 |
| 1A4 | low | 7 | 7 | 16 | M | NA | 0 | NA |
| 1B1 | high | 1 | NA | 4 | NA | NA | 0 | 3 |
| 1B2 | high | 3 | 7 | 18 | M | NA | 0 | 15 |
| 1B3 | high | 5 | NA | 21 | F | 3.539 | 0 | 16 |
| 1B4 | high | 7 | 7 | 27 | M | NA | 0 | NA |
| 1C1 | control | NA | 7 | 25 | F | 3.344 | 0 | NA |
| 1C2 | control | NA | 7 | 14 | NA | NA | 0 | NA |
| 1C3 | control | NA | 7 | 15 | F | 3.079 | 0 | NA |
| 1C4 | control | NA | 7 | 25 | F | 3.297 | 0 | NA |
| 20A1 | low | 1 | 7 | 22 | M | NA | 130000 | 21 |
| 20A2 | low | 3 | 7 | 20 | F | 2.95 | 150000 | 17 |
| 20A3 | low | 5 | 7 | 20 | M | NA | 10000 | 15 |
| 20A4 | low | 7 | 7 | 22 | M | NA | 0 | NA |
| 20B1 | high | 1 | 7 | 21 | M | NA | 1570000 | 20 |
| 20B2 | high | 3 | 7 | 26 | F | NA | 720000 | 23 |
| 20B3 | high | 5 | 6 | 31 | M | NA | 130000 | 26 |
| 20B4 | high | 7 | 7 | 21 | F | 3.3385 | 0 | NA |
| 20C1 | control | NA | 7 | 21 | F | 2.993 | 0 | NA |
| 20C2 | control | NA | 7 | 10 | F | 3.214 | 0 | NA |
| 20C3 | control | NA | 9 | 10 | NA | NA | 0 | NA |
| 20C4 | control | NA | 7 | 12 | F | 3.52 | 0 | NA |
| 21A1 | low | 1 | 7 | 9 | F | NA | 0 | 8 |
| 21A2 | low | 3 | 7 | 17 | F | 3.1375 | 60000 | 14 |
| 21A3 | low | 5 | 7 | 23 | F | 3.186 | 190000 | 18 |
| 21A4 | low | 7 | 9 | 29 | F | 3.309 | 0 | NA |
| 21B1 | high | 1 | 8 | 21 | F | 3.45 | 380000 | 20 |
| 21B2 | high | 3 | 7 | 22 | F | 3.359 | 0 | 19 |
| 21B3 | high | 5 | 7 | 24 | F | 3.487 | 40000 | 19 |
| 21B4 | high | 7 | 8 | 27 | F | 3.173 | 160000 | NA |

| | | | | | | | | |
|---|---|---|---|---|---|---|---|---|
| 21C1 | control | NA | 7 | 29 | M | NA | 0 | NA |
| 21C2 | control | NA | 7 | 8 | NA | NA | 0 | NA |
| 21C3 | control | NA | 7 | 29 | M | NA | 0 | NA |
| 21C4 | control | NA | 7 | 25 | F | 3.687 | 0 | NA |
| 22A1 | low | 1 | 7 | 15 | M | NA | 0 | 14 |
| 22A2 | low | 3 | 6 | 29 | M | NA | 750000 | 26 |
| 22A3 | low | 5 | 7 | 9 | NA | NA | 0 | 4 |
| 22A4 | low | 7 | 7 | 22 | M | NA | 0 | NA |
| 22B1 | high | 1 | 7 | 21 | F | NA | 2770000 | 20 |
| 22B2 | high | 3 | 8 | 26 | F | 3.433 | 400000 | 23 |
| 22B3 | high | 5 | 7 | 20 | M | NA | 20000 | 15 |
| 22B4 | high | 7 | 7 | 27 | F | 3.396 | 0 | NA |
| 22C1 | control | NA | 7 | 22 | M | NA | 0 | NA |
| 22C2 | control | NA | 8 | 36 | F | NA | 0 | NA |
| 22C3 | control | NA | 8 | 26 | M | NA | 0 | NA |
| 22C4 | control | NA | 7 | 26 | M | NA | 0 | NA |
| 23A1 | low | 1 | 7 | 14 | F | 3.697 | 100000 | 13 |
| 23A2 | low | 3 | 7 | 10 | M | NA | 0 | 7 |
| 23A3 | low | 5 | 7 | 16 | M | NA | 0 | 11 |
| 23A4 | low | 7 | 7 | 25 | F | 3.334 | 0 | NA |
| 23B1 | high | 1 | 7 | 23 | M | NA | 30000 | 22 |
| 23B2 | high | 3 | 7 | 22 | F | 3.437 | 140000 | 19 |
| 23B3 | high | 5 | 7 | 21 | M | NA | 50000 | 16 |
| 23B4 | high | 7 | 7 | 26 | F | 3.335 | 0 | NA |
| 23C1 | control | NA | 8 | 21 | M | NA | 0 | NA |
| 23C2 | control | NA | 7 | 27 | F | NA | 0 | NA |
| 23C3 | control | NA | 7 | 28 | F | NA | 0 | NA |
| 23C4 | control | NA | 7 | 26 | F | 3.45 | 0 | NA |
| 24A1 | low | 1 | 7 | 26 | M | NA | 980000 | 25 |
| 24A2 | low | 3 | 7 | 23 | M | NA | 160000 | 20 |
| 24A3 | low | 5 | 8 | 25 | F | 3.24 | 400000 | 20 |
| 24A4 | low | 7 | 7 | 29 | F | 3.391 | 0 | NA |
| 24B1 | high | 1 | 7 | 22 | M | NA | 720000 | 21 |
| 24B2 | high | 3 | 7 | 26 | F | 3.31 | 400000 | 23 |
| 24B3 | high | 5 | 7 | 23 | M | NA | 170000 | 18 |
| 24B4 | high | 7 | 7 | 29 | M | 3.388 | 0 | NA |
| 24C1 | control | NA | 8 | 26 | F | 3.389 | 0 | NA |
| 24C2 | control | NA | 7 | 20 | F | 3.2595 | 0 | NA |
| 24C3 | control | NA | 8 | 21 | M | NA | 0 | NA |
| 24C4 | control | NA | 7 | 22 | NA | NA | 0 | NA |
| 25A1 | low | 1 | 7 | 19 | F | 3.497 | 90000 | 18 |
| 25A2 | low | 3 | 8 | 32 | F | 3.181 | 1930000 | 29 |
| 25A3 | low | 5 | 7 | 10 | NA | NA | 0 | 5 |
| 25A4 | low | 7 | 9 | 35 | M | NA | 90000 | NA |
| 25B1 | high | 1 | 7 | 12 | F | 3.313 | 10000 | 11 |
| 25B2 | high | 3 | 8 | 22 | F | 3.2 | 690000 | 19 |
| 25B3 | high | 5 | NA | 7 | NA | NA | 0 | 2 |
| 25B4 | high | 7 | 7 | 21 | M | NA | 0 | NA |
| 25C1 | control | NA | 9 | 32 | M | NA | 0 | NA |
| 25C2 | control | NA | 8 | 33 | F | 3.35 | 0 | NA |
| 25C3 | control | NA | 7 | 24 | M | NA | 0 | NA |

| | | | | | | | | | |
|---|---|---|---|---|---|---|---|---|---|
| 25C4 | control | NA | | 7 | 27 | M | NA | 0 | NA |
| 26A1 | low | | 1 | 8 | 18 | F | 3.407 | 2430000 | 17 |
| 26A2 | low | | 3 | 7 | 19 | F | 3.644 | 50000 | 16 |
| 26A3 | low | | 5 | 7 | 19 | F | 3.512 | 30000 | 14 |
| 26A4 | low | | 7 | 7 | 10 | F | 3.374 | 0 | NA |
| 26B1 | high | | 1 | NA | 6 | NA | NA | 0 | 5 |
| 26B2 | high | | 3 | 7 | 24 | F | 3.202 | 90000 | 21 |
| 26B3 | high | | 5 | 7 | 29 | M | NA | 540000 | 24 |
| 26B4 | high | | 7 | 7 | 32 | F | 3.059 | 0 | NA |
| 26C1 | control | NA | | NA | 6 | NA | NA | 0 | NA |
| 26C2 | control | NA | | 7 | 28 | F | 3.02 | 0 | NA |
| 26C3 | control | NA | | 8 | 21 | F | 3.567 | 0 | NA |
| 26C4 | control | NA | | 7 | 10 | F | 3.106 | 0 | NA |
| 27A1 | low | | 1 | 8 | 11 | M | NA | 560000 | 10 |
| 27A2 | low | | 3 | 7 | 20 | M | NA | 50000 | 17 |
| 27A3 | low | | 5 | 7 | 26 | F | 3.437 | 0 | 21 |
| 27A4 | low | | 7 | 7 | 22 | F | 3.188 | 0 | NA |
| 27B1 | high | | 1 | 7 | 21 | F | 3.697 | 1240000 | 20 |
| 27B2 | high | | 3 | 7 | 23 | M | NA | 30000 | 20 |
| 27B3 | high | | 5 | 7 | 20 | F | 3.389 | 30000 | 15 |
| 27B4 | high | | 7 | 8 | 25 | M | NA | 220000 | NA |
| 27C1 | control | NA | | 7 | 23 | F | NA | 0 | NA |
| 27C2 | control | NA | | 7 | 10 | F | NA | 0 | NA |
| 27C3 | control | NA | | 7 | 10 | M | NA | 0 | NA |
| 27C4 | control | NA | | 7 | 33 | M | NA | 0 | NA |
| 28A1 | low | | 1 | 7 | 18 | F | 3.4475 | 640000 | 17 |
| 28A2 | low | | 3 | 8 | 21 | F | 3.3175 | 40000 | 18 |
| 28A3 | low | | 5 | 7 | 17 | M | NA | 0 | 12 |
| 28A4 | low | | 7 | 7 | 22 | F | 2.75 | 0 | NA |
| 28B1 | high | | 1 | 7 | 26 | M | NA | 750000 | 25 |
| 28B2 | high | | 3 | 8 | 28 | M | NA | 200000 | 25 |
| 28B3 | high | | 5 | 7 | 23 | F | 2.32 | 0 | 18 |
| 28B4 | high | | 7 | 7 | 10 | F | 3.23 | 0 | NA |
| 28C1 | control | NA | | 7 | 22 | F | 3.468 | 0 | NA |
| 28C2 | control | NA | | 7 | 24 | F | 3.24 | 0 | NA |
| 28C3 | control | NA | | 8 | 10 | NA | NA | 0 | NA |
| 28C4 | control | NA | | 7 | 21 | M | NA | 0 | NA |
| 29A1 | low | | 1 | 6 | 23 | M | NA | 280000 | 22 |
| 29A2 | low | | 3 | 7 | 17 | F | NA | 0 | 14 |
| 29A3 | low | | 5 | 8 | 9 | NA | NA | 0 | 4 |
| 29A4 | low | | 7 | 7 | 13 | M | NA | 0 | NA |
| 29B1 | high | | 1 | 7 | 14 | M | NA | 200000 | 13 |
| 29B2 | high | | 3 | 7 | 10 | M | NA | 0 | 7 |
| 29B3 | high | | 5 | 7 | 22 | F | 3.3 | 150000 | 17 |
| 29B4 | high | | 7 | 7 | 29 | M | NA | 0 | NA |
| 29C1 | control | NA | | 8 | 13 | F | 3.575 | 0 | NA |
| 29C2 | control | NA | | 7 | 26 | F | 3.229 | 0 | NA |
| 29C3 | control | NA | | 7 | 21 | F | NA | 0 | NA |
| 29C4 | control | NA | | 7 | 33 | M | NA | 0 | NA |
| 2A1 | low | | 1 | NA | 4 | NA | NA | 0 | 3 |
| 2A2 | low | | 3 | 7 | 13 | F | 3.452 | 0 | 10 |

| | | | | | | | | |
|---|---|---|---|---|---|---|---|---|
| 2A3 | low | 5 | 7 | 26 | F | 3.407 | 20000 | 21 |
| 2A4 | low | 7 | 7 | 22 | M | NA | 0 | NA |
| 2B1 | high | 1 | 7 | 20 | F | 3.48 | 290000 | 19 |
| 2B2 | high | 3 | 7 | 25 | F | 3.32 | 185000 | 22 |
| 2B3 | high | 5 | 7 | 25 | M | NA | 50000 | 20 |
| 2B4 | high | 7 | 7 | 25 | M | NA | 0 | NA |
| 2C1 | control | NA | 7 | 16 | M | NA | 0 | NA |
| 2C2 | control | NA | 7 | 15 | F | 3.251 | 0 | NA |
| 2C3 | control | NA | 7 | 20 | M | NA | 0 | NA |
| 2C4 | control | NA | 7 | 13 | M | NA | 0 | NA |
| 30A1 | low | 1 | 7 | 13 | F | 3.692 | 0 | 12 |
| 30A2 | low | 3 | NA | 8 | NA | NA | 0 | 6 |
| 30A3 | low | 5 | 7 | 13 | M | NA | 0 | 8 |
| 30A4 | low | 7 | 7 | 13 | M | NA | 0 | NA |
| 30B1 | high | 1 | NA | 4 | NA | NA | 0 | 3 |
| 30B2 | high | 3 | 7 | 14 | M | NA | 0 | 11 |
| 30B3 | high | 5 | 7 | 21 | M | NA | 120000 | 16 |
| 30B4 | high | 7 | NA | 5 | NA | NA | 0 | NA |
| 30C1 | control | NA | NA | 6 | NA | NA | 0 | NA |
| 30C2 | control | NA | 7 | 21 | F | 3.417 | 0 | NA |
| 30C3 | control | NA | NA | 7 | NA | NA | 0 | NA |
| 30C4 | control | NA | 9 | 16 | NA | NA | 0 | NA |
| 31A1 | low | 1 | 7 | 12 | M | NA | 0 | 11 |
| 31A2 | low | 3 | 7 | 15 | M | NA | 40000 | 12 |
| 31A3 | low | 5 | 7 | 21 | F | NA | 100000 | 16 |
| 31A4 | low | 7 | 8 | 25 | F | 3.498 | 20000 | NA |
| 31B1 | high | 1 | 7 | 25 | F | 3.16 | 1120000 | 24 |
| 31B2 | high | 3 | 8 | 13 | F | 3.465 | 0 | 10 |
| 31B3 | high | 5 | 7 | 23 | M | NA | 180000 | 18 |
| 31B4 | high | 7 | 7 | 12 | NA | NA | 0 | NA |
| 31C1 | control | NA | 7 | 22 | F | 3.188 | 0 | NA |
| 31C2 | control | NA | 7 | 19 | F | 3.526 | 0 | NA |
| 31C3 | control | NA | 7 | 22 | F | 3.13 | 0 | NA |
| 31C4 | control | NA | 7 | 23 | F | 3.33 | 0 | NA |
| 32A1 | low | 1 | 7 | 23 | M | NA | 130000 | 22 |
| 32A2 | low | 3 | 7 | 9 | M | NA | 0 | 6 |
| 32A3 | low | 5 | 7 | 33 | M | NA | 20000 | 28 |
| 32A4 | low | 7 | 7 | 23 | M | NA | 0 | NA |
| 32B1 | high | 1 | 7 | 13 | F | 3.667 | 60000 | 12 |
| 32B2 | high | 3 | 7 | 20 | F | 3.609 | 540000 | 17 |
| 32B3 | high | 5 | 7 | 22 | M | 3.343 | 380000 | 17 |
| 32B4 | high | 7 | 7 | 27 | F | 3.176 | 0 | NA |
| 32C1 | control | NA | 8 | 34 | F | 3.156 | 0 | NA |
| 32C2 | control | NA | 7 | 25 | F | 3.623 | 0 | NA |
| 32C3 | control | NA | 8 | 33 | F | 3.344 | 0 | NA |
| 32C4 | control | NA | 7 | 25 | F | 3.18 | 0 | NA |
| 33A1 | low | 1 | 7 | 16 | M | NA | 320000 | 15 |
| 33A2 | low | 3 | 7 | 30 | M | NA | 310000 | 27 |
| 33A3 | low | 5 | 7 | 26 | M | NA | 20000 | 21 |
| 33A4 | low | 7 | 7 | 21 | M | NA | 0 | NA |
| 33B1 | high | 1 | NA | 4 | NA | NA | 0 | 3 |

| | | | | | | | | |
|---|---|---|---|---|---|---|---|---|
| 33B2 | high | 3 | 7 | 16 | F | 3.371 | 0 | 13 |
| 33B3 | high | 5 | 7 | 25 | M | NA | 20000 | 20 |
| 33B4 | high | 7 | 8 | 22 | M | NA | 0 | NA |
| 33C1 | control | NA | 7 | 33 | F | 3.042 | 0 | NA |
| 33C2 | control | NA | 8 | 26 | F | 3.467 | 0 | NA |
| 33C3 | control | NA | 7 | 20 | M | NA | 0 | NA |
| 33C4 | control | NA | 7 | 20 | F | 3.579 | 0 | NA |
| 34A1 | low | 1 | 7 | 22 | F | 3.52 | 580000 | 21 |
| 34A2 | low | 3 | 7 | 27 | F | 3.296 | 350000 | 24 |
| 34A3 | low | 5 | 7 | 25 | F | 3.242 | 150000 | 20 |
| 34A4 | low | 7 | 7 | 22 | F | 2.75 | 0 | NA |
| 34B1 | high | 1 | NA | 4 | NA | NA | 0 | 3 |
| 34B2 | high | 3 | 7 | 22 | F | 3.185 | 100000 | 19 |
| 34B3 | high | 5 | 7 | 26 | F | 3.162 | 210000 | 21 |
| 34B4 | high | 7 | 7 | 23 | F | 3.364 | 0 | NA |
| 34C1 | control | NA | 7 | 24 | M | 3.363 | 0 | NA |
| 34C2 | control | NA | 7 | 26 | M | NA | 0 | NA |
| 34C3 | control | NA | 7 | 21 | F | 3.799 | 0 | NA |
| 34C4 | control | NA | 7 | 27 | F | 3.61 | 0 | NA |
| 35A1 | low | 1 | 7 | 20 | F | 3.57 | 710000 | 19 |
| 35A2 | low | 3 | 7 | 15 | F | 3.225 | 0 | 12 |
| 35A3 | low | 5 | 7 | 25 | F | 3.297 | 100000 | 20 |
| 35A4 | low | 7 | 7 | 20 | F | 3.535 | 0 | NA |
| 35B1 | high | 1 | 7 | 23 | M | NA | 90000 | 22 |
| 35B2 | high | 3 | 7 | 23 | M | NA | 50000 | 20 |
| 35B3 | high | 5 | 7 | 21 | F | NA | 40000 | 16 |
| 35B4 | high | 7 | 7 | 23 | M | NA | 0 | NA |
| 35C1 | control | NA | 7 | 9 | NA | NA | 0 | NA |
| 35C2 | control | NA | 7 | 12 | M | NA | 0 | NA |
| 35C3 | control | NA | 7 | 15 | F | 3.497 | 0 | NA |
| 35C4 | control | NA | 7 | 15 | F | 3.402 | 0 | NA |
| 36A1 | low | 1 | 7 | 14 | F | 3.52 | 80000 | 13 |
| 36A2 | low | 3 | 7 | 15 | M | NA | 0 | 12 |
| 36A3 | low | 5 | 7 | 8 | NA | NA | 0 | 3 |
| 36A4 | low | 7 | 7 | 30 | M | NA | 0 | NA |
| 36B1 | high | 1 | NA | 4 | NA | NA | 0 | 3 |
| 36B2 | high | 3 | 7 | 10 | F | 3.272 | 0 | 7 |
| 36B3 | high | 5 | 7 | 26 | M | NA | 70000 | 21 |
| 36B4 | high | 7 | 7 | 26 | F | 3.331 | 0 | NA |
| 36C1 | control | NA | 7 | 34 | F | 2.85 | 0 | NA |
| 36C2 | control | NA | 7 | 10 | M | NA | 0 | NA |
| 36C3 | control | NA | 7 | 21 | M | NA | 0 | NA |
| 36C4 | control | NA | NA | 5 | NA | NA | 0 | NA |
| 37A1 | low | 1 | 7 | 19 | F | 3.294 | 1130000 | 18 |
| 37A2 | low | 3 | 7 | 17 | M | NA | 550000 | 14 |
| 37A3 | low | 5 | NA | 8 | NA | NA | 0 | 3 |
| 37A4 | low | 7 | 7 | 35 | F | 3.687 | 40000 | NA |
| 37B1 | high | 1 | 7 | 21 | M | NA | 0 | 20 |
| 37B2 | high | 3 | 8 | 19 | M | NA | 380000 | 16 |
| 37B3 | high | 5 | 7 | 28 | F | 3.28 | 270000 | 23 |
| 37B4 | high | 7 | 8 | 27 | F | 3.4 | 68000 | NA |

| | | | | | | | | |
|---|---|---|---|---|---|---|---|---|
| 37C1 | control | NA | 8 | 10 | F | 3.2 | 0 | NA |
| 37C2 | control | NA | 8 | 23 | M | NA | 0 | NA |
| 37C3 | control | NA | 7 | 23 | F | 3.444 | 0 | NA |
| 37C4 | control | NA | 8 | 14 | F | 3.366 | 0 | NA |
| 38A1 | low | 1 | 7 | 32 | M | NA | 670000 | 31 |
| 38A2 | low | 3 | 7 | 22 | F | 3.464 | 30000 | 19 |
| 38A3 | low | 5 | 7 | 11 | F | 3.523 | 0 | 6 |
| 38A4 | low | 7 | 7 | 28 | M | NA | 0 | NA |
| 38B1 | high | 1 | 7 | 15 | F | NA | 50000 | 14 |
| 38B2 | high | 3 | 7 | 24 | F | 3.006 | 210000 | 21 |
| 38B3 | high | 5 | 7 | 16 | F | 3.415 | 30000 | 11 |
| 38B4 | high | 7 | 7 | 14 | F | NA | 0 | NA |
| 38C1 | control | NA | 7 | 14 | F | 3.462 | 0 | NA |
| 38C2 | control | NA | 8 | 23 | M | NA | 0 | NA |
| 38C3 | control | NA | 7 | 13 | NA | NA | 0 | NA |
| 38C4 | control | NA | 7 | 26 | M | NA | 0 | NA |
| 39A1 | low | 1 | 7 | 26 | F | 3.382 | 1920000 | 25 |
| 39A2 | low | 3 | 7 | 30 | M | NA | 250000 | 27 |
| 39A3 | low | 5 | 7 | 25 | F | 3.24 | 300000 | 20 |
| 39A4 | low | 7 | 8 | 23 | F | 3.34 | 0 | NA |
| 39B1 | high | 1 | 8 | 28 | F | 3.566 | 90000 | 27 |
| 39B2 | high | 3 | 8 | 23 | M | NA | 670000 | 20 |
| 39B3 | high | 5 | 8 | 22 | F | 3.217 | 10000 | 17 |
| 39B4 | high | 7 | 8 | 34 | M | NA | 0 | NA |
| 39C1 | control | NA | 7 | 21 | F | 3.566 | 0 | NA |
| 39C2 | control | NA | 7 | 26 | F | 3.36 | 0 | NA |
| 39C3 | control | NA | 8 | 10 | F | 3.6 | 0 | NA |
| 39C4 | control | NA | 7 | 16 | M | NA | 0 | NA |
| 3A1 | low | 1 | 7 | 26 | M | NA | 1210000 | 25 |
| 3A2 | low | 3 | 7 | 22 | M | NA | 0 | 19 |
| 3A3 | low | 5 | 8 | 34 | M | NA | 610000 | 29 |
| 3A4 | low | 7 | 8 | 25 | M | NA | 0 | NA |
| 3B1 | high | 1 | 7 | 21 | M | NA | 300000 | 20 |
| 3B2 | high | 3 | 7 | 21 | M | NA | 70000 | 18 |
| 3B3 | high | 5 | 8 | 26 | M | NA | 230000 | 21 |
| 3B4 | high | 7 | 7 | 21 | M | NA | 0 | NA |
| 3C1 | control | NA | 7 | 20 | F | 3.278 | 0 | NA |
| 3C2 | control | NA | 8 | 26 | F | 3.43 | 0 | NA |
| 3C3 | control | NA | 8 | 20 | F | 3.313 | 0 | NA |
| 3C4 | control | NA | 7 | 21 | M | NA | 0 | NA |
| 40A1 | low | 1 | 7 | 19 | F | 3.596 | 1600000 | 18 |
| 40A2 | low | 3 | 7 | 19 | M | NA | 200000 | 16 |
| 40A3 | low | 5 | 7 | 21 | M | NA | 0 | 16 |
| 40A4 | low | 7 | 7 | 10 | M | NA | 0 | NA |
| 40B1 | high | 1 | 7 | 21 | F | 3.569 | 210000 | 20 |
| 40B2 | high | 3 | 8 | 13 | M | NA | 0 | 10 |
| 40B3 | high | 5 | 7 | 23 | F | 3.443 | 800000 | 18 |
| 40B4 | high | 7 | 7 | 29 | F | 3.208 | 0 | NA |
| 40C1 | control | NA | 6 | 36 | F | 3.132 | 0 | NA |
| 40C2 | control | NA | 7 | 29 | M | NA | 0 | NA |
| 40C3 | control | NA | 7 | 27 | F | 3.349 | 0 | NA |

| | | | | | | | | |
|---|---|---|---|---|---|---|---|---|
| 40C4 | control | NA | 7 | 23 | F | 3.498 | 0 | NA |
| 41A1 | low | 1 | 7 | 23 | F | 3.66 | 80000 | 22 |
| 41A2 | low | 3 | 7 | 12 | F | NA | 0 | 9 |
| 41A3 | low | 5 | 7 | 20 | F | 3.075 | 210000 | 15 |
| 41A4 | low | 7 | 7 | 23 | M | NA | 0 | NA |
| 41B1 | high | 1 | 7 | 10 | M | NA | 0 | 9 |
| 41B2 | high | 3 | 8 | 15 | F | 3.128 | 0 | 12 |
| 41B3 | high | 5 | 8 | 31 | M | NA | 870000 | 26 |
| 41B4 | high | 7 | 7 | 25 | M | NA | 0 | NA |
| 41C1 | control | NA | 7 | 33 | F | 3.423 | 0 | NA |
| 41C2 | control | NA | 7 | 20 | M | NA | 0 | NA |
| 41C3 | control | NA | 7 | 10 | F | 3.3325 | 0 | NA |
| 41C4 | control | NA | 7 | 25 | M | NA | 0 | NA |
| 42A1 | low | 1 | 7 | 22 | F | 3.465 | 60000 | 21 |
| 42A2 | low | 3 | 7 | 14 | M | NA | 0 | 11 |
| 42A3 | low | 5 | 7 | 14 | F | 3.202 | 0 | 9 |
| 42A4 | low | 7 | 7 | 29 | M | NA | 0 | NA |
| 42B1 | high | 1 | 7 | 21 | M | NA | 410000 | 20 |
| 42B2 | high | 3 | 8 | 9 | NA | NA | 0 | 6 |
| 42B3 | high | 5 | 9 | 28 | F | NA | 420000 | 23 |
| 42B4 | high | 7 | 7 | 9 | NA | NA | 0 | NA |
| 42C1 | control | NA | 7 | 25 | F | 3.37 | 0 | NA |
| 42C2 | control | NA | 7 | 13 | F | 3.243 | 0 | NA |
| 42C3 | control | NA | 8 | 14 | F | 3.42 | 0 | NA |
| 42C4 | control | NA | NA | 5 | NA | NA | 0 | NA |
| 43A1 | low | 1 | 7 | 21 | F | 3.32 | 680000 | 20 |
| 43A2 | low | 3 | 7 | 10 | NA | NA | 0 | 7 |
| 43A3 | low | 5 | 7 | 9 | NA | NA | 0 | 4 |
| 43A4 | low | 7 | 7 | 15 | M | NA | 0 | NA |
| 43B1 | high | 1 | 7 | 15 | F | 3.363 | 270000 | 14 |
| 43B2 | high | 3 | 7 | 21 | F | 3.316 | 90000 | 18 |
| 43B3 | high | 5 | 7 | 16 | F | 3.541 | 0 | 11 |
| 43B4 | high | 7 | 7 | 15 | F | 3.356 | 0 | NA |
| 43C1 | control | NA | 7 | 13 | M | NA | 0 | NA |
| 43C2 | control | NA | 7 | 24 | M | NA | 0 | NA |
| 43C3 | control | NA | 7 | 26 | F | 3.572 | 0 | NA |
| 43C4 | control | NA | 7 | 21 | F | NA | 0 | NA |
| 44A1 | low | 1 | NA | 7 | NA | NA | 0 | NA |
| 44A2 | low | 3 | 7 | 23 | F | 3.142 | 160000 | 20 |
| 44A3 | low | 5 | 7 | 24 | F | 3.398 | 32000 | 19 |
| 44A4 | low | 7 | 8 | 19 | M | NA | 0 | 12 |
| 44B1 | high | 1 | 7 | 21 | M | NA | 250000 | 20 |
| 44B2 | high | 3 | 8 | 23 | F | 2.856 | 60000 | 20 |
| 44B3 | high | 5 | 8 | 20 | F | 3.488 | 10000 | 15 |
| 44B4 | high | 7 | 7 | 25 | F | 3.419 | 0 | NA |
| 44C1 | control | NA | 7 | 26 | M | NA | 0 | NA |
| 44C2 | control | NA | 7 | 13 | F | 3.475 | 0 | NA |
| 44C3 | control | NA | 7 | 22 | M | NA | 0 | NA |
| 44C4 | control | NA | 7 | 24 | F | 3.153 | 0 | NA |
| 45A1 | low | 1 | 7 | 23 | M | NA | 920000 | 22 |
| 45A2 | low | 3 | 7 | 26 | F | 3.18 | 2570000 | 23 |

| | | | | | | | | |
|---|---|---|---|---|---|---|---|---|
| 45A3 | low | 5 | 7 | 10 | M | NA | 0 | 5 |
| 45A4 | low | 7 | 8 | 25 | F | 3.579 | 1050000 | 18 |
| 45B1 | high | 1 | NA | 4 | NA | NA | 0 | 3 |
| 45B2 | high | 3 | 7 | 11 | M | NA | 0 | 10 |
| 45B3 | high | 5 | 7 | 16 | NA | NA | 0 | 11 |
| 45B4 | high | 7 | 7 | 10 | NA | NA | 0 | NA |
| 45C1 | control | NA | 7 | 26 | M | NA | 0 | NA |
| 45C2 | control | NA | 7 | 11 | F | 3.473 | 0 | NA |
| 45C3 | control | NA | 7 | 29 | M | NA | 0 | NA |
| 45C4 | control | NA | 7 | 22 | F | 2.963 | 0 | NA |
| 46A1 | low | 1 | 8 | 19 | F | NA | 1310000 | 18 |
| 46A2 | low | 3 | 8 | 27 | F | NA | 180000 | 24 |
| 46A3 | low | 5 | NA | 5 | NA | NA | 0 | 0 |
| 46A4 | low | 7 | 7 | 23 | F | 3.521 | 0 | NA |
| 46B1 | high | 1 | NA | 4 | NA | NA | 0 | 3 |
| 46B2 | high | 3 | NA | 4 | NA | NA | 0 | 1 |
| 46B3 | high | 5 | 7 | 21 | F | 3.602 | 40000 | 16 |
| 46B4 | high | 7 | 7 | 28 | NA | 3.498 | 0 | NA |
| 46C1 | control | NA | 7 | 23 | M | NA | 0 | NA |
| 46C2 | control | NA | 8 | 33 | M | NA | 0 | NA |
| 46C3 | control | NA | NA | 5 | NA | 3.663 | 0 | NA |
| 46C4 | control | NA | 7 | 16 | F | NA | 0 | NA |
| 47A1 | low | 1 | 7 | 12 | M | NA | 0 | 11 |
| 47A2 | low | 3 | 7 | 19 | M | NA | 40000 | 16 |
| 47A3 | low | 5 | 9 | 14 | M | NA | 0 | 9 |
| 47A4 | low | 7 | 7 | 21 | M | NA | 0 | NA |
| 47B1 | high | 1 | 8 | 14 | F | 3.343 | 560000 | 13 |
| 47B2 | high | 3 | 9 | 16 | M | NA | 30000 | 13 |
| 47B3 | high | 5 | 8 | 9 | F | NA | 0 | 3 |
| 47B4 | high | 7 | 8 | 26 | F | 3.388 | 30000 | NA |
| 47C1 | control | NA | 8 | 26 | M | NA | 0 | NA |
| 47C2 | control | NA | 9 | 29 | M | NA | 0 | NA |
| 47C3 | control | NA | 8 | 21 | M | NA | 0 | NA |
| 47C4 | control | NA | 8 | 13 | F | 3.53 | 0 | NA |
| 48A1 | low | 1 | 7 | 9 | NA | NA | 0 | 8 |
| 48A2 | low | 3 | 7 | 15 | F | 3.339 | 760000 | 13 |
| 48A3 | low | 5 | 7 | 9 | NA | NA | 0 | 4 |
| 48A4 | low | 7 | 7 | 28 | NA | NA | 0 | NA |
| 48B1 | high | 1 | 7 | 15 | F | 3.543 | 0 | 14 |
| 48B2 | high | 3 | NA | 6 | NA | NA | 0 | 3 |
| 48B3 | high | 5 | 7 | 9 | NA | NA | 0 | 4 |
| 48B4 | high | 7 | 7 | 8 | NA | NA | 0 | NA |
| 48C1 | control | NA | 7 | 26 | F | 3.227 | 0 | NA |
| 48C2 | control | NA | 7 | 9 | NA | NA | 0 | NA |
| 48C3 | control | NA | 7 | 13 | F | 3.609 | 0 | NA |
| 48C4 | control | NA | 7 | 6 | NA | NA | 0 | NA |
| 49A1 | low | 1 | 7 | 10 | M | NA | 0 | 9 |
| 49A2 | low | 3 | 7 | 21 | F | 3.279 | 140000 | 18 |
| 49A3 | low | 5 | 7 | 21 | F | 3.598 | 10000 | 16 |
| 49A4 | low | 7 | 7 | 23 | F | 3.187 | 0 | NA |
| 49B1 | high | 1 | 7 | 23 | F | NA | 340000 | 22 |

| | | | | | | | | |
|---|---|---|---|---|---|---|---|---|
| 49B2 | high | 3 | 8 | 19 | M | NA | 0 | 16 |
| 49B3 | high | 5 | NA | 6 | NA | NA | 0 | 1 |
| 49B4 | high | 7 | 7 | 22 | F | 3.479 | 0 | NA |
| 49C1 | control | NA | 7 | 22 | F | 3.208 | 0 | NA |
| 49C2 | control | NA | 7 | 34 | F | 3.298 | 0 | NA |
| 49C3 | control | NA | 8 | 15 | F | 3.4735 | 0 | NA |
| 49C4 | control | NA | 7 | 22 | F | 3.254 | 0 | NA |
| 4A1 | low | 1 | 7 | 11 | M | NA | 0 | 10 |
| 4A2 | low | 3 | 7 | 26 | M | NA | 380000 | 23 |
| 4A3 | low | 5 | 7 | 18 | F | 3.578 | 0 | 13 |
| 4A4 | low | 7 | 8 | 20 | F | 3.45 | 0 | NA |
| 4B1 | high | 1 | 7 | 23 | F | 3.525 | 1870000 | 22 |
| 4B2 | high | 3 | 7 | 29 | F | 3.085 | 700000 | 26 |
| 4B3 | high | 5 | 7 | 26 | F | 3.526 | 20000 | 21 |
| 4B4 | high | 7 | 8 | 26 | F | 3.503 | 20000 | NA |
| 4C1 | control | NA | 7 | 34 | M | NA | 0 | NA |
| 4C2 | control | NA | 8 | 12 | M | NA | 0 | NA |
| 4C3 | control | NA | 8 | 10 | M | NA | 0 | NA |
| 4C4 | control | NA | 7 | 25 | F | 3.408 | 0 | NA |
| 50A1 | low | 1 | 7 | 19 | F | 3.29 | 60000 | 18 |
| 50A2 | low | 3 | 7 | 25 | M | NA | 380000 | 22 |
| 50A3 | low | 5 | 7 | 22 | F | 2.95 | 0 | 17 |
| 50A4 | low | 7 | 7 | 19 | F | 3.298 | 0 | NA |
| 50B1 | high | 1 | 7 | 19 | F | 3.593 | 420000 | 18 |
| 50B2 | high | 3 | 7 | 21 | F | NA | 350000 | 18 |
| 50B3 | high | 5 | 7 | 10 | M | NA | 0 | 5 |
| 50B4 | high | 7 | 8 | 26 | M | NA | 140000 | NA |
| 50C1 | control | NA | 7 | 18 | M | NA | 0 | NA |
| 50C2 | control | NA | 7 | 23 | M | NA | 0 | NA |
| 50C3 | control | NA | 8 | 22 | M | NA | 0 | NA |
| 50C4 | control | NA | 7 | 26 | M | NA | 0 | NA |
| 5A1 | low | 1 | 7 | 21 | M | NA | 0 | 20 |
| 5A2 | low | 3 | 7 | 26 | F | 3.216 | 0 | 23 |
| 5A3 | low | 5 | 8 | 22 | M | NA | 0 | 17 |
| 5A4 | low | 7 | 7 | 25 | M | NA | 0 | NA |
| 5B1 | high | 1 | 7 | 19 | F | 3.353 | 2190000 | 18 |
| 5B2 | high | 3 | 7 | 19 | M | 3.196 | 290000 | 16 |
| 5B3 | high | 5 | 8 | 24 | F | NA | 30000 | 19 |
| 5B4 | high | 7 | 7 | 21 | M | NA | 0 | NA |
| 5C1 | control | NA | 7 | 13 | M | NA | 0 | NA |
| 5C2 | control | NA | 8 | 23 | M | NA | 0 | NA |
| 5C3 | control | NA | NA | 6 | NA | NA | 0 | NA |
| 5C4 | control | NA | 8 | 26 | F | 3.343 | 0 | NA |
| 6A1 | low | 1 | 7 | 22 | M | NA | 0 | 21 |
| 6A2 | low | 3 | 8 | 23 | F | 3.31 | 340000 | 20 |
| 6A3 | low | 5 | 8 | 24 | M | NA | 70000 | 19 |
| 6A4 | low | 7 | 8 | 26 | F | 3.613 | 0 | NA |
| 6B1 | high | 1 | 7 | 22 | F | 3.456 | 1890000 | 21 |
| 6B2 | high | 3 | 8 | 17 | M | NA | 20000 | 14 |
| 6B3 | high | 5 | 7 | 22 | F | 3.282 | 120000 | 17 |
| 6B4 | high | 7 | 8 | 9 | F | 3.196 | 0 | NA |

| | | | | | | | | |
|---|---|---|---|---|---|---|---|---|
| 6C1 | control | NA | 8 | 20 | F | 3.392 | 0 | NA |
| 6C2 | control | NA | 8 | 30 | F | 3.572 | 0 | NA |
| 6C3 | control | NA | 8 | 22 | M | NA | 0 | NA |
| 6C4 | control | NA | 8 | 26 | F | 3.34 | 0 | NA |
| 7A1 | low | 1 | 7 | 16 | M | NA | 50000 | 15 |
| 7A2 | low | 3 | 7 | 23 | M | NA | 720000 | 20 |
| 7A3 | low | 5 | 7 | 15 | F | NA | 0 | 10 |
| 7A4 | low | 7 | 7 | 27 | F | 3.485 | 0 | NA |
| 7B1 | high | 1 | 7 | 25 | M | NA | 640000 | 24 |
| 7B2 | high | 3 | 7 | 22 | F | 3.4 | 170000 | 19 |
| 7B3 | high | 5 | 8 | 26 | F | 3.252 | 0 | 21 |
| 7B4 | high | 7 | 7 | 26 | F | 3.345 | 0 | NA |
| 7C1 | control | NA | 7 | 10 | M | NA | 0 | NA |
| 7C2 | control | NA | 9 | 10 | NA | 3.3913 | 0 | NA |
| 7C3 | control | NA | 7 | 19 | F | 3.643 | 0 | NA |
| 7C4 | control | NA | 7 | 24 | M | NA | 0 | NA |
| 8A1 | low | 1 | 8 | 20 | F | 3.647 | 1220000 | 19 |
| 8A2 | low | 3 | 7 | 10 | NA | NA | 0 | 7 |
| 8A3 | low | 5 | 8 | 26 | F | 3.09 | 20000 | 21 |
| 8A4 | low | 7 | 7 | 17 | F | 3.398 | 0 | NA |
| 8B1 | high | 1 | NA | 4 | NA | NA | 0 | 3 |
| 8B2 | high | 3 | 7 | 30 | F | 3.572 | 270000 | 27 |
| 8B3 | high | 5 | 6 | 14 | F | 3.285 | 0 | 9 |
| 8B4 | high | 7 | 7 | 21 | F | 3.503 | 0 | NA |
| 8C1 | control | NA | 7 | 25 | F | 3.125 | 0 | NA |
| 8C2 | control | NA | 7 | 24 | F | 3.223 | 0 | NA |
| 8C3 | control | NA | 7 | 24 | M | NA | 0 | NA |
| 8C4 | control | NA | 8 | 14 | F | 3.207 | 0 | NA |
| 9A1 | low | 1 | 7 | 12 | F | NA | 20000 | 11 |
| 9A2 | low | 3 | 7 | 25 | F | 3.288 | 2700000 | 22 |
| 9A3 | low | 5 | 8 | 19 | F | 3.675 | 150000 | NA |
| 9A4 | low | 7 | 7 | 26 | NA | NA | 0 | NA |
| 9B1 | high | 1 | NA | 4 | NA | NA | 0 | 3 |
| 9B2 | high | 3 | 8 | 9 | F | 3.388 | 0 | 6 |
| 9B3 | high | 5 | 7 | 26 | M | NA | 90000 | 21 |
| 9B4 | high | 7 | 7 | 21 | M | NA | 0 | NA |
| 9C1 | control | NA | 9 | 11 | F | 3.045 | 0 | NA |
| 9C2 | control | NA | 7 | 16 | F | 3.521 | 0 | NA |
| 9C3 | control | NA | 7 | 22 | F | NA | 0 | NA |
| 9C4 | control | NA | NA | 10 | NA | NA | 0 | NA |
